# RNA localization mechanisms transcend cell morphology

**DOI:** 10.1101/2022.04.14.488401

**Authors:** Raeann Goering, Ankita Arora, J. Matthew Taliaferro

## Abstract

RNA molecules are localized to specific subcellular regions through interactions between RNA regulatory elements and RNA binding proteins (RBPs). Generally, our knowledge of the mechanistic details behind the localization of a given RNA is restricted to a particular cell type. Here, we show that RNA/RBP interactions that regulate RNA localization in one cell type predictably regulate localization in other cell types with vastly different morphologies. To determine transcriptome-wide RNA spatial distributions across the apicobasal axis of human intestinal epithelial cells, we used our recently developed RNA proximity labeling technique, Halo-seq. We found that mRNAs encoding ribosomal proteins (RP mRNAs) were strongly localized to the basal pole of these cells. Using reporter transcripts and single molecule RNA FISH, we found that pyrimidine-rich TOP motifs in the 5′ UTRs of RP mRNAs were sufficient to drive basal RNA localization. Interestingly, the same TOP motifs were also sufficient to drive RNA localization to the neurites of mouse neuronal cells. In both cell types, the regulatory activity of the TOP motif was dependent on it being at the extreme 5′ end of the transcript, was abolished upon perturbation of the TOP-binding protein LARP1, and was reduced upon inhibition of kinesins. To extend these findings, we compared subcellular RNAseq data from neuronal and epithelial cells. We found that the basal compartment of epithelial cells and the projections of neuronal cells were enriched for highly similar sets of RNAs, indicating that broadly similar mechanisms may be transporting RNAs in both cell types. These findings identify the first RNA element known to regulate RNA localization across the apicobasal axis of epithelial cells, establish LARP1 as an RNA localization regulator, and demonstrate that RNA localization mechanisms cut across cell morphologies.

## INTRODUCTION

The post-transcriptional regulation of RNA allows for the fine tuning of the expression of genetic information. These post-transcriptional processes, including alternative splicing, translational regulation, RNA decay, and subcellular RNA localization, are often regulated by RNA binding proteins (RBPs) that exert their function upon their RNA targets by recognizing and binding specific sequence motifs. Because the binding of RNA by RBPs is largely governed by specific sequence-based interactions, RBPs often bind similar RNA targets across cell types. Therefore, RBPs often exert their regulatory function upon the same RNA targets, regardless of cellular context (Matia-González et al., 2015; Van Nostrand et al., 2020).

Many RBPs are ubiquitously expressed across tissues and generally elicit the same post-transcriptional regulatory behaviors on their target RNAs in many cell types (Gerstberger et al., 2014). The subcellular localization of an RNA, though, is at least conceptually tightly linked to cell morphology. If the interaction between an RBP and an RNA sequence in neurons result in a transcript’s localization to axons, where will the same transcript localize in a cell type that doesn’t have axons yet nevertheless expresses the same regulatory RBP? (**Figure 1A**) Can we define spatial equivalents for RNA destinations across cells with vastly different morphologies?

**Figure 1.**
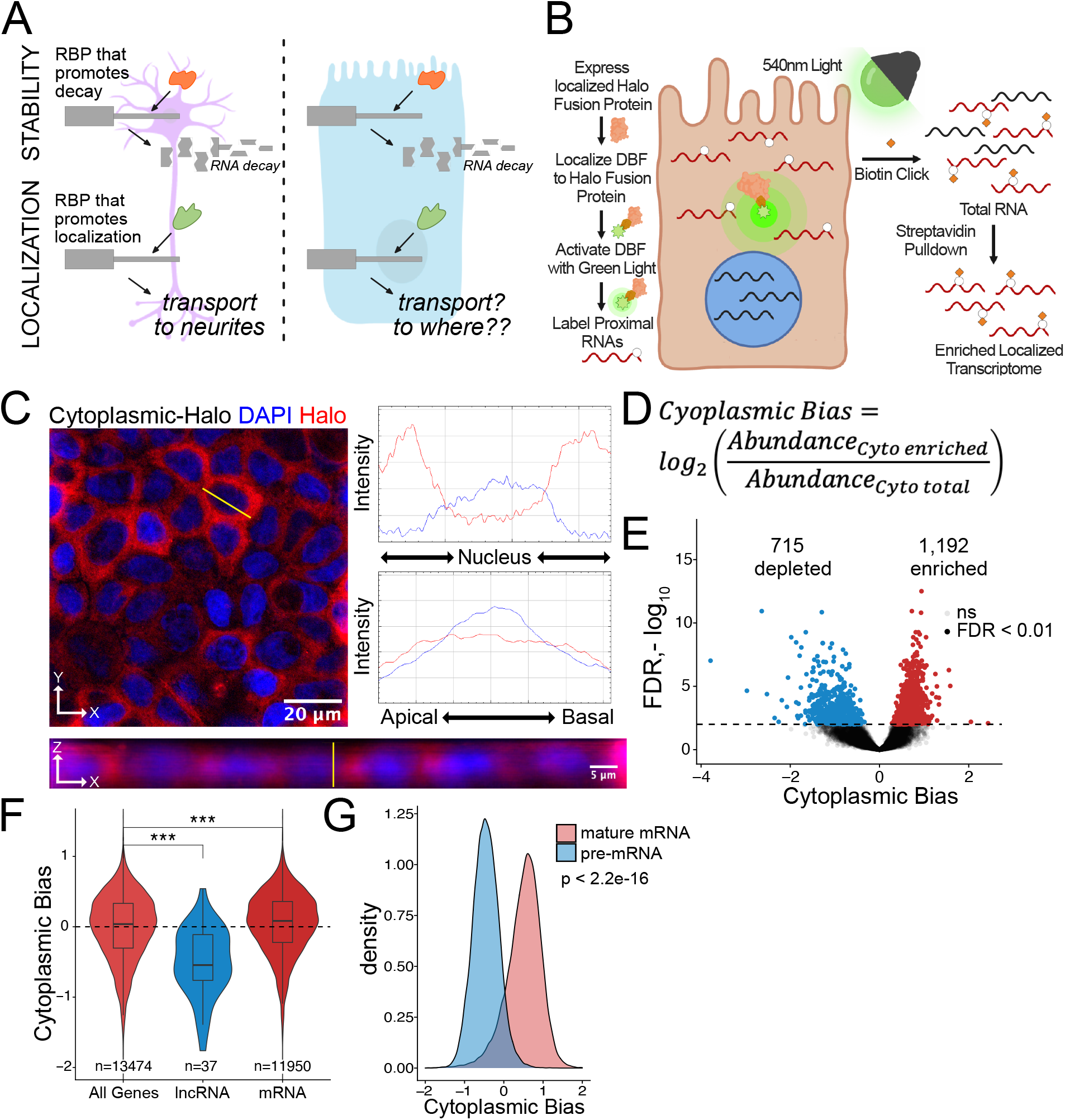
Halo-seq enriches cytoplasmic localized RNA molecules in C2bbe1 monolayers. (A) RNA regulatory processes are generally driven through the interaction of an RBP and an RNA sequence element. Because RBPs are often widely expressed, many of these processes (exemplified here by RNA decay) operate across cellular contexts. The orange RBP binds a particular sequence within a transcript to promote its degradation in both cell types. However, because RNA localization is intimately linked with cell morphology, if an RBP/RNA interaction promotes RNA localization in one cell type, how the same interaction would affect RNA localization in another cell type is unclear. (B) Overview of the Halo-seq method. A HaloTag is genetically fused to a protein with specific localization. DBF-conjugated Halo ligands are specifically recruited to the Halo fusion proteins due to their high affinity and covalent binding to HaloTag protein domains. When exposed to green light, DBF produces oxygen radicals that, in combination with propargylamine, alkynylate DBF-proximal RNAs. “Click” chemistry covalently attaches biotin moieties to alkynylated RNAs *in vitro*, facilitating their enrichment via streptavidin pulldown, thereby enriching localized RNAs for high-throughput sequencing. (C) HaloTag protein fusions to P65 are localized to the cytoplasm. HaloTag fusions are visualized through the addition of a Halo ligand fluorophore shown in red. DAPI stain marks the nuclei in blue. Profile lines across the XY image demonstrate exclusion of Halo signal from the nucleus and in the XZ-projected image demonstrate no apical or basal bias. Images are an average projection through the XZ axis of many cells. (D) Equation for Cytoplasmic Bias. Cytoplasmic RNA localization is calculated with a metric termed Cytoplasmic Bias as the log2 of a transcript’s abundance in the streptavidin enriched fraction divided by its abundance in the input total RNA. (E) Genes with differing Cytoplasmic Bias following Halo-seq RNA labeling using the cytoplasmic-Halo fusion. FDR calculated by DESeq2 (F) Cytoplasmic Bias for defined classes of RNAs. P-values were calculated using a Wilcoxon rank-sum test. (G) Cytoplasmic Bias for unspliced, intron-containing pre-mRNAs and spliced mature mRNAs. Cytoplasmic Bias was calculated for the spliced and unspliced isoform of each transcript. P-values were calculated using a Wilcoxon rank-sum test. ns (not significant) represents p > 0.05, * p < 0.05, ** p < 0.01, *** p < 0.001 and **** represents p < 0.0001.

For a given RNA, much of what is known about the regulation of its localization, especially in terms of sequence motifs and the RBPs that bind them, has been elucidated in a single cell type. Many examples of the molecular rules that govern RNA transport to neuronal projections (Arora et al., 2021; Goering et al., 2020; von Kügelgen et al., 2021; Mikl et al., 2021; Zappulo et al., 2017) or across the posterior/anterior axis of the Drosophila embryos (Ephrussi and Lehmann, 1992; Hachet and Ephrussi, 2004; Jambor et al., 2014; Lécuyer et al., 2007; Lipshitz and Smibert, 2000) have been defined. However, how these RBPs and RNA elements would regulate RNA localization in cell types or morphologies outside of those that they were discovered in is generally unknown.

RNA localization is widely studied in neuronal models as their long projections are amenable to both microscopy and subcellular fractionation. Hundreds of RNAs are actively transported into neuronal projections (Cajigas et al., 2012; Gumy et al., 2011; Taliaferro et al., 2016; Zappulo et al., 2017; Zivraj et al., 2010). Many of these RNAs are not neuron specific and are expressed widely throughout many other tissues (von Kügelgen and Chekulaeva, 2020; Zappulo et al., 2017). Again, whether these RNA are localized in other cell types is largely unknown.

Intestinal enterocytes line the intestine as a single cell monolayer. Enterocytes are polarized across their apicobasal axis with specialized regions that perform specific functions (Klunder et al., 2017). The apical regions of enterocytes are responsible for nutrient absorption from the gut while the basal regions export processed nutrients to the lymph and bloodstream (Chen et al., 2018; Goodman, 2010). Many proteins and RNAs are asymmetrically localized across the apicobasal axis to aid in these spatially distinct functions, but little is known about the RNA elements and RBPs that regulate a transcript’s location within epithelial cells (Moor et al., 2017; Ryder and Lerit, 2018).

Using transcriptome-wide assays, we reveal that the neurites of neurons and the basal pole of epithelial cells are enriched for similar RNAs. We further find that specific RNA regulatory elements and RBPs that regulate RNA localization in one cell type can predictably do so in other cell types with very different morphologies as well as in different species. These results hint at conserved mechanisms of RNA localization and potentially a predictable, underlying code for RNA localization compatible with vastly different subcellular locations and cellular morphologies.

## RESULTS

### Halo-seq allows for isolation of spatially distinct RNA populations from C2bbe1 monolayers

C2bbe1 cells, a subclone of the human CaCo-2 cell line, recapitulate human intestinal enterocytes by spontaneously creating polarized monolayers with differentiated brush-borders in culture (Peterson and Mooseker, 1992). For easy genetic manipulation, we created a C2bbe1 cell line with a blasticidin resistance gene surrounded by *loxP* recombination sites in the AAVS1 safe harbor locus. This system allows for the controlled insertion of doxycycline-inducible transgenes using *cre* recombinase (Khandelia et al., 2011). This knock-in cell line was created with CRISPR/Cas9, an sgRNA targeting the AAVS1 locus, and a homologous repair donor plasmid containing the *loxP* cassette (**Figure S1A**). Following selection with blasticidin, integration of the cassette was confirmed by PCR (**Figure S1B**). This cell line is amenable to quick and efficient *cre*mediated cassette switching allowing for stable doxycycline-inducible expression of any RNA or protein of interest (**Figure S1C**).

We used our recently developed proximity labeling technique Halo-seq (Engel et al., 2021) to assay RNA localization across the apicobasal axis of C2bbe1 monolayers (**Figure 1B**). Halo-seq relies on the expression of a Halo-tagged protein with specific localization to a subcellular compartment of interest. HaloTags are small protein domains that covalently bind to small molecules known as Halo ligands (Los et al., 2008). Through conjugation to a Halo ligand backbone, the photo-activatable singlet oxygen generator dibromofluorescein (DBF) can be specifically recruited to the Halo-tagged protein and therefore share the same subcellular distribution as the fusion protein.

When exposed to green light, DBF generates oxygen radicals that are restricted to a radius of approximately 100 nm around the Halo fusion protein (Jin et al., 1995; Li et al., 2018). RNA bases within this cloud are oxidized by the radicals, making them susceptible to nucleophilic attack by propargylamine, a cell permeable alkyne (Ali et al., 2004; Li et al., 2018). Halo-fusion-proximal RNAs are therefore selectively alkynylated *in situ*.

Following total RNA isolation, alkynylated transcripts are biotinylated using Cu(I)-catalyzed azide-alkyne cycloaddition (CuACC) “Click” chemistry with biotin azide (Hein et al., 2008). Halo-proximal transcripts are therefore selectively biotinylated by this reaction allowing for purification of localized RNAs from total RNA via streptavidin pulldowns. By comparing the abundances of transcripts in total RNA and streptavidin-purified RNA samples using high-throughput sequencing, transcripts that were Halo-proximal can be identified.

To first assay the efficiency with which Halo-seq can enrich localized RNAs in C2bbe1 monolayers, we targeted the cytoplasmic compartment for analysis with Halo-seq. Because differences between the cytoplasmic and nuclear transcriptomes are well characterized (Engel et al., 2021; Zaghlool et al., 2021), we reasoned that this would provide a good benchmark of the technique in C2bbe1 monolayers. We fused a HaloTag domain to a cytoplasmically restricted NF-kappa B subunit, P65. This doxycycline-inducible transgene was integrated into the genome of C2bbe1 cells using cre/*loxP* recombination. The doxycycline-inducible expression and size of the cytoplasmic-Halo fusion were confirmed by gel electrophoresis with fluorescent Halo ligands (**Figure S1D**). We also confirmed the cytoplasmic localization of the Halo fusion protein *in situ* using fluorescent Halo ligands (**Figure 1C**). We then visualized the site-specific alkynylation of biomolecules following DBF-mediated oxidation by performing azide-alkyne cycloaddition *in situ* with Cy3-conjugated azides. The presence of Cy3 labeled molecules required the addition of DBF, indicating that the observed alkynylation was due to the Halo-seq procedure (**Figure S1E**). These experiments revealed that both the cytoplasmic-Halo fusion and the molecules it labeled were restricted to the cytoplasm, indicating that the DBF-mediated labeling was spatially specific.

To determine if we could biotinylate alkynated RNA, we performed Halo-seq in differentiated C2bbe1 monolayers expressing the cytoplasmic-Halo. Following in-cell alkynylation and *in vitro* Click-mediated biotinylation, we visualized biotinylated RNA using RNA dot blots with streptavidin-HRP. We observed strong RNA biotinylation signal that was dependent upon the addition of DBF to cells (**Figure S1F**). We then incubated biotinylated RNA samples from DBF-treated and untreated cells with streptavidin-coated beads. For three DBF-treated samples, we recovered an average of 1.6% of the total RNA sample from the streptavidin-coated beads. In contrast, we only recovered an average of 0.05% of the total RNA sample from DBF-untreated samples, further illustrating that RNA biotinylation requires DBF treatment (**Figure S1G**).

We then created rRNA-depleted RNAseq libraries from three streptavidin input and enriched RNA samples from DBF-treated cells. Following high-throughput sequencing, we quantified transcript and gene abundances in each sample using salmon and tximport (Patro et al., 2017; Soneson et al., 2015). As expected, samples separated primarily by fraction (input vs. streptavidin pulldown) in a principal component analysis (PCA) plot of transcript expression profiles (**Figure S1H**).

To quantify cytoplasmic RNA localization, we created a metric called cytoplasmic bias (CB), which we defined as the log2 of a transcript’s abundance in the streptavidin-pulldown samples compared to its abundance in the input RNA samples (**Figure 1D**). We used DESeq2 to identify genes that were significantly enriched in streptavidin-pulldown samples compared to input samples (Love et al., 2014). These represent genes whose transcripts were enriched in the cytoplasm. Using this approach, 1,192 genes were identified as significantly cytoplasmically enriched (FDR < 0.01) while 715 genes were significantly depleted and therefore likely enriched in the nucleus (**Figure 1E, Table S1**).

We then looked at CB values for lncRNAs specifically. As a class, these transcripts are known to be enriched in the nucleus (Lubelsky and Ulitsky, 2018; Shukla et al., 2018), and therefore should be depleted from the streptavidin-pulldown samples. As expected, lncRNA transcripts had significantly lower CB values, indicating that Halo-seq was faithfully reporting on RNA localization in C2bbe1 cells (**Figure 1F**). Additionally, we used a custom salmon index to simultaneously quantify intron-containing and fully spliced versions of every transcript. Spliced transcripts are exported to the cytoplasm while intron-containing transcripts are often restricted to the nucleus (Luo and Reed, 1999). Further validating Halo-seq, we observed that intron-containing pre-mRNA transcripts generally had negative CB values, indicating their nuclear enrichment. Conversely, intron-lacking, mature transcripts of the same genes had positive CB values, indicating their cytoplasmic enrichment (**Figure 1G**). These results provided confidence that Halo-seq identifies localized RNAs in C2bbe1 monolayers.

### Halo-seq identifies RNAs differentially localized across the apicobasal axis of C2bbe1 monolayers

To identify RNAs localized across the apicobasal axis of enterocytes, we first needed to ensure that we could establish polarized monolayers of C2bbe1 cells. We plated C2bbe1 cells on porous transwell inserts at high confluency and allowed spontaneous differentiation to occur for 7 days (Masuda et al., 2011; Peterson and Mooseker, 1992). This produced highly polarized monolayers with the expected spatially distinct localization of the polarity protein markers Ezrin and the sodium-potassium pump across the apicobasal axis, suggesting mature enterocyte-like organization (**Figure S2A**).

To interrogate apicobasal RNA localization using Halo-seq, we created HaloTag fusion proteins that specifically localize to either the apical or basal poles of differentiated cells. In intestinal epithelial cells, podocalyxin-like protein 1 (PODXL) is apically localized while a subunit of the sodium-potassium pump (ATP1A1) is basally localized (Deborde et al., 2008; Meder et al., 2005). Using these membrane proteins, we created transgenes where HaloTags were fused to their N-termini such that the HaloTag domain was inside the cell. These doxycycline-inducible transgenes were integrated into the C2bbe1 genome using the cre/*loxP* strategy described in Figure S1C.

We confirmed doxycycline-inducible expression of the full length HaloTag fusion proteins using gel electrophoresis and fluorescent Halo ligands (**Figure S2B**). The subcellular localization of both HaloTag fusions was then assessed using fluorescent Halo ligands. The PODXL fusion (henceforth referred to as apical-Halo) was restricted to the apical region of cells while the ATP1A1 fusion (henceforth referred to as basal-Halo) was restricted to the basal pole (**Figure 2A-B**). To assess the localization of alkynylated molecules produced by the Halo-seq protocol with these fusions, we visualized alkynylated molecules using *in situ* Click chemistry with a fluorescent azide as in Figure S1E. We found that in cells expressing apical-Halo, alkynylated molecules were concentrated toward the apical poles of cells. Conversely, in cells expressing Basal-Halo, alkynylated molecules were concentrated toward the basal pole. In both cases, alkynylation was dependent upon the addition of DBF to cells (**Figure S2C-D**). These results indicate that Halo-seq can spatially specifically alkynylate biomolecules across the apicobasal axis in epithelial monolayers, giving us confidence in the ability of Halo-seq to isolate RNA molecules that were localized to the apical and basal poles of epithelial cells.

**Figure 2.**
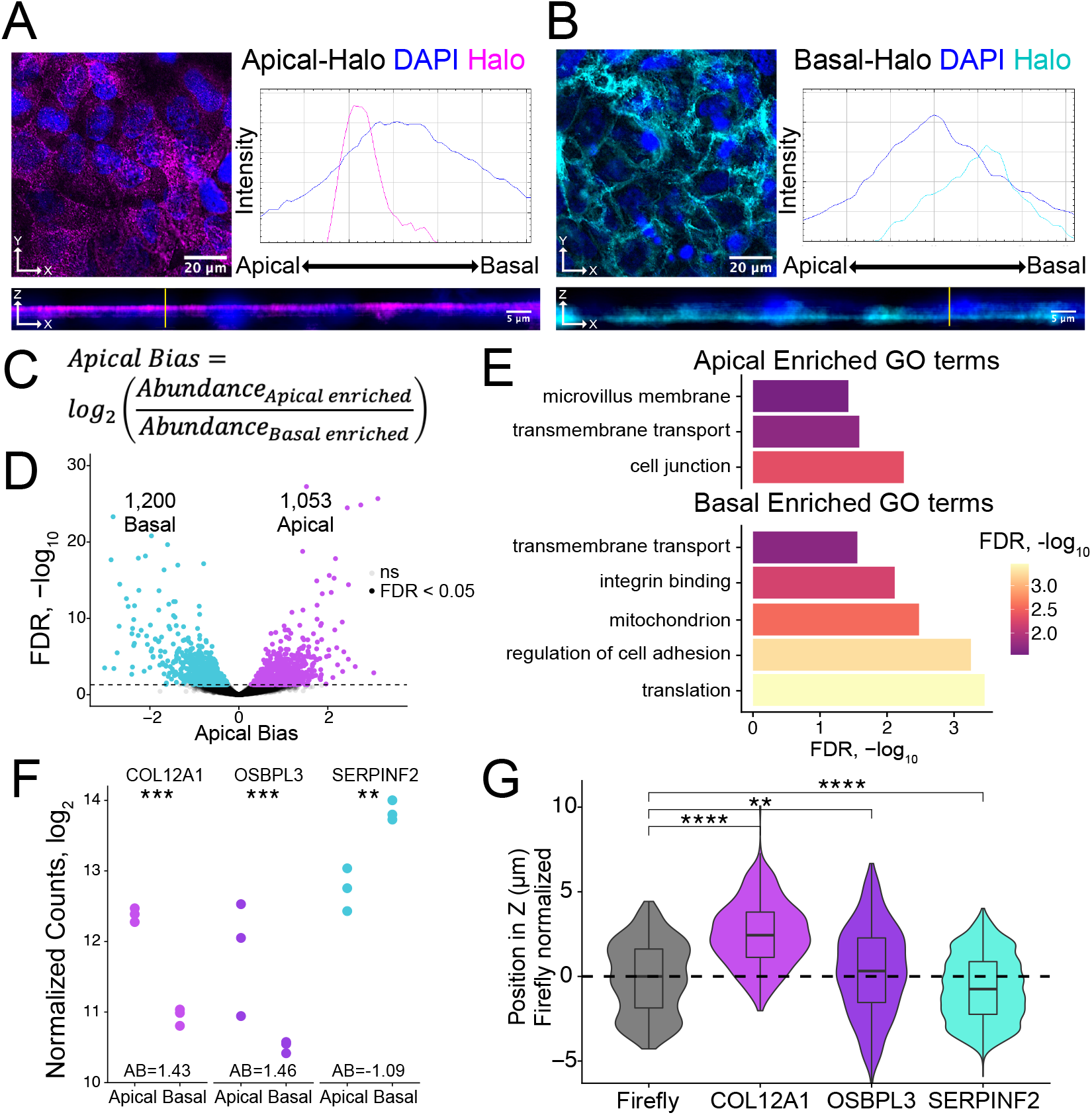
Halo-seq enriches RNAs at the apical and basal poles of enterocyte monolayers. (A) HaloTag protein fusions to PODXL and (B) ATP1A1 are localized to the apical and basal compartments respectively. HaloTag domains are visualized through the addition of a Halo ligand fluorophore shown in magenta (apical) and cyan (basal). DAPI stain marks the nuclei in blue. Profile lines across the XZ image demonstrate their apical or basal bias. Images are an average projection through the XZ axis of many cells. (C) Equation for Apical Bias. RNA localization across the apicobasal axis is calculated with a metric termed Apical Bias as the log2 of a transcript’s abundance in the apical enriched RNA fraction divided by its abundance in the basal enriched fraction. (D) Genes with differing Apical Bias following Halo-seq RNA labeling using the apical and basal-Halo fusions. FDR values calculated by DESeq2. (E) Enriched gene ontology (GO) terms associated with proteins encoded by RNAs identified as localized to the apical or basal pole of enterocytes. FDR calculated by topGO. (F) DESeq2 calculated normalized counts of candidate apical (COL12A1 and OSBPL3) and basal (SERPINF2) localized RNAs in the apical and basal enriched sequencing samples. P-values calculated by DESeq2. (G) smFISH puncta position in Z. Firefly luciferase RNA included as a non-localized control. P-values were calculated using a Wilcoxon rank-sum test. ns (not significant) represents p > 0.05, * p < 0.05, ** p < 0.01, *** p < 0.001 and **** represents p < 0.0001.

We then confirmed the of the apical-Halo and basal-Halo fusions to result in RNA biotinylation using RNA dot blots (**Figure S2E**) and purified the resulting products using streptavidin-coated beads. The apical-Halo and basal-Halo fusion proteins produced less biotinylated RNA than the cytoplasmic-Halo fusion. However, for both the apical and basal fusions, we recovered significantly more RNA following streptavidin pulldown from cells treated with DBF than from control untreated cells (**Figure S2F**), further indicating the ability of these Halo fusions to mediate selective RNA biotinylation.

We isolated apically and basally localized RNA by performing Halo-seq in triplicate with apical-Halo and basal-Halo. We then quantified the resulting RNA samples using high-throughput sequencing of rRNA-depleted RNAseq libraries, calculating transcript and gene abundances using salmon and tximport (Patro et al., 2017; Soneson et al., 2015). Sequencing replicates of all Halo-seq samples first separate by genotype (PC1xPC2) then by fraction (PC2xPC3) in a PCA of gene expression profiles (**Figure S2G**). To identify RNAs that were differentially localized across the apicobasal axis, we calculated an apical bias (AB) value for each gene. We defined this metric as the log2 ratio of a gene’s abundance in the apical-Halo pulldown samples compared to its abundance in the basal-Halo pulldown samples (**Figure 2C, Table S1**). Genes with positive AB values are therefore apically-enriched while those with negative AB values are basally-enriched. AB values and their associated p values were calculated using DESeq2 (Love et al., 2014). With this approach, transcripts from 1,053 genes were found to be significantly apically localized (FDR < 0.05) while transcripts from 1,200 genes were significantly basally localized (**Figure 2D**). These results suggest that many RNAs are asymmetrically distributed across the apicobasal axis in epithelial cells.

To preliminarily characterize the apical and basal transcriptomes, we used TopGO (Alexa et al., 2010) to identify enriched gene ontology (GO) terms associated with the proteins encoded by RNAs localized to each compartment (**Figure 2E**). The term “trans-membrane transport” was significantly associated with RNAs localized to both compartments (p < 0.05), consistent with the fact that both the apical and basal membranes of enterocytes play major roles in nutrient import and export (Kiela and Ghishan, 2016; Snoeck et al., 2005). Unique GO terms associated with apically localized RNAs included “cell junctions” including apical tight junctions, and “microvillus membrane” which is exclusively associated with enterocyte apical membrane (Crawley et al., 2014; Sluysmans et al., 2017). Proteins encoded by basally localized RNAs were enriched (p < 0.05) for functions associated with cell adhesion and integrin binding, both of which are important for basal membrane anchoring to the extracellular matrix (Yurchenco, 2011).

RNAs from nuclear-encoded mitochondrial proteins were also found to be basally enriched. Because these RNAs are often localized to mitochondria due to their translation on the mitochondrial surface (Corral-Debrinski M. et al., 2000; Williams et al., 2014), this would imply that mitochondria themselves may also be basally localized in C2bbe1 monolayers. To test this, we visualized mitochondria using both immunofluorescence and a mitochondrial-specific dye, mitoView. Both methods indicated that the localization of mitochondria was, in fact, basally biased (**Figure S2H**). These results gave us confidence in the reliability of the Halo-seq quantifications.

### Halo-seq results are consistent with previous RNA localization datasets in polarized enterocytes

A previous study used laser capture microdissection (LCM) and RNAseq to assess RNA localization across the apicobasal axis in adult mouse intestine tissue slices (Moor et al., 2017). In this study, RNA samples from apical and basal LCM slices were compared to define RNAs differentially distributed across the apicobasal axis of these cells. To assess the quality of our Halo-seq data, we compared our AB values to those defined in this dataset. Using the mouse intestine data, we calculated AB values for all genes and compared them to Halo-seq derived AB values of their human orthologs. We found a modest (R = 0.08), yet positive and significant (p = 0.003) correlation between these two datasets (**Figure S2I**). This correlation overcomes differences in samples (cultured cells vs. tissue), methodology (Halo-seq vs. LCM), and species (human vs. mouse). We found this correlation to be encouraging and further suggestive of the ability of Halo-seq to faithfully report on RNA localization across the apicobasal axis in epithelial cells.

### Validation of Halo-seq results using single molecule RNA FISH

To experimentally validate Halo-seq’s ability to identify transcripts differentially localized across the apicobasal axis, we performed single molecule fluorescent *in situ* hybridization (smFISH) to visualize endogenous transcripts within C2bbe1 monolayers (Tsanov et al., 2016). We selected two transcripts with positive AB values, *COL12A1* and *OSBPL3*, and one transcript with a negative AB value, *SERPINF2*, for validation (**Figure 2F, S2J**). Spots corresponding to RNA molecules in smFISH images were detected using FISH-quant (Mueller et al., 2013; Tsanov et al., 2016) and the 3D coordinates of each spot were identified. This allowed calculation of the position of each transcript’s position along the image’s Z axis, which corresponds to the cell’s apicobasal axis. When analyzing smFISH images for each endogenous transcript, we ensured that the number of spots detected per cell was proportional to the expression levels of the transcript as calculated by RNAseq (**Figure S2K-L**), suggesting that each custom designed smFISH probe set (Tsanov et al., 2016) was faithfully reporting on the location of its cognate transcript. By comparing to the localization of an exogenous, nonlocalized transcript encoding Firefly luciferase as a nonlocalized control, we found that *COL12A1* and *OSBPL3* transcripts were significantly apically localized, while *SERPINF2* transcripts were basally localized (**Figure 2G, S2M**). These results are consistent with their AB values as determined by Halo-seq, giving us additional confidence in the reliability of the Halo-seq dataset.

### 5′ TOP motifs are sufficient to drive RNA transcripts to the basal pole of epithelial monolayers

Using our Halo-seq derived dataset of RNAs localized across the apicobasal axis of enterocytes, we observed that mRNAs encoding ribosomal proteins (RP mRNAs) were, as a class, strongly basally localized (**Figure 3A**). This is consistent with previous observations made in adult mouse intestines (Moor et al., 2017), as well as with a LCM-based study of RNA localization in *Drosophila* follicular epithelial cells (Cassella and Ephrussi, 2021) (**Figure S3A**), further suggesting that our model and technique are faithfully recapitulating phenomena that occur *in vivo*. While RP mRNAs are known to be basally localized (Cassella and Ephrussi, 2021; Moor et al., 2017), the mechanisms underlying this localization are unknown.

**Figure 3.**
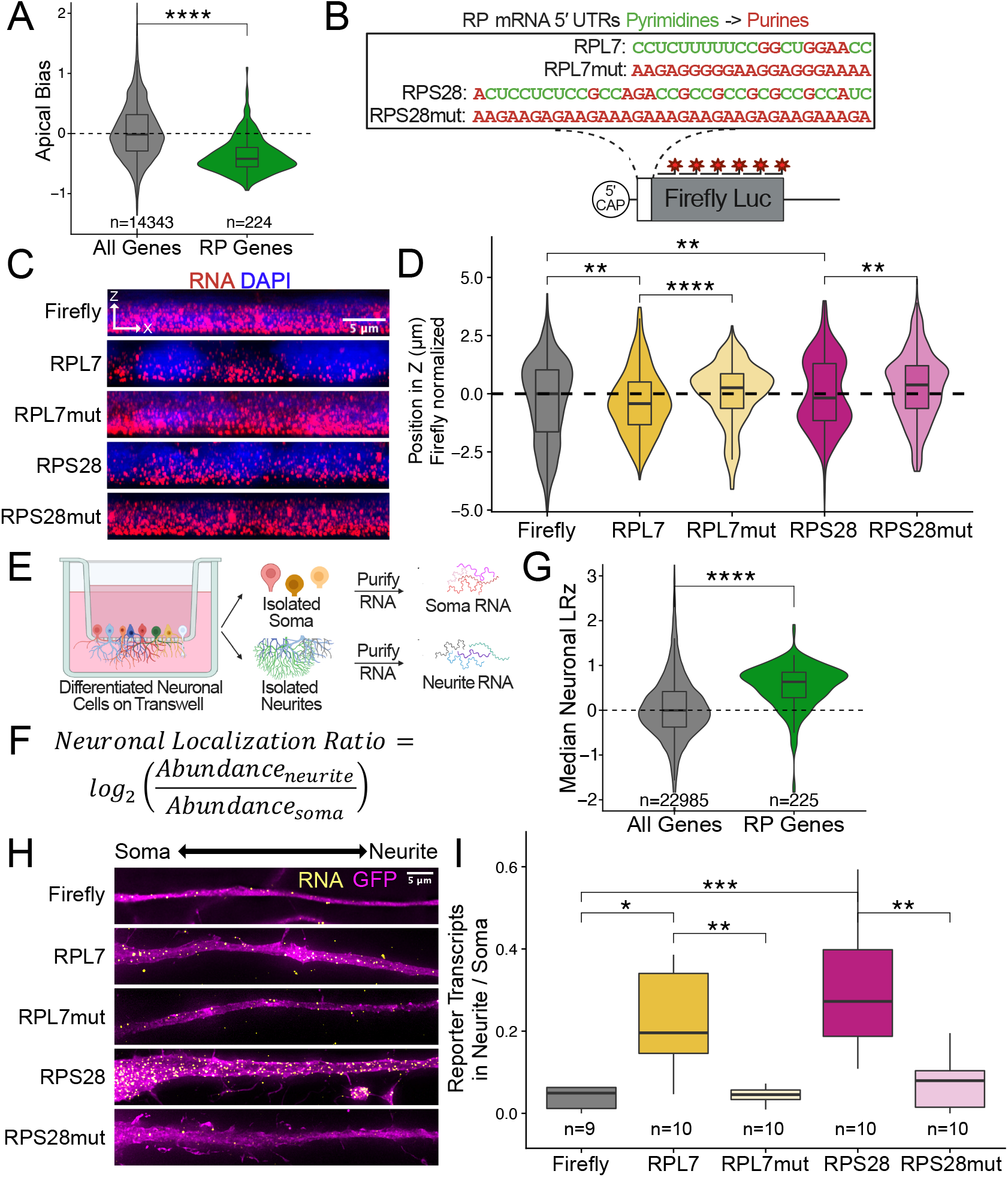
Ribosomal protein mRNAs are localized in a 5′ TOP dependent manner in many cell types. Apical Bias for ribosomal protein (RP) genes as compared to all genes. P-values were calculated using a Wilcoxon rank-sum test. (B) Schematic of ribosomal protein 5′ TOP reporter constructs.Mutant versions were created by swapping each pyrimidine for a purine. (C) Representative images of smFISH spots for each 5′ TOP reporter construct. RNA signal shown in red while DAPI stains mark the nuclei blue. Images are a max projection through the XZ axis of many cells. (D) smFISH puncta position in Z normalized to median untagged Firefly luciferase transcript position. P-values were calculated using a Wilcoxon rank-sum test. (E) Schematic of neuronal subcellular fractionation and localized RNA isolation. (F) Equation for Neuronal Localization Ratio. Neuronal RNA localization is calculated with a metric termed Neuronal Localization Ratio as the log2 of a gene’s abundance in the neurite fraction divided by its abundance in the soma fraction. (G) Neuronal Localization Ratio (LR) for ribosomal protein genes as compared to all genes. P-values were calculated using a Wilcoxon rank-sum test. (H) Representative images of smFISH for each 5′ TOP reporter construct. Images are a max projection through the XY axis of a single neurite positioned with the soma to the left. RNA signal shown in yellow while cell outlines marked by GFP signal shown as magenta. (I) Number of smFISH puncta in neurites normalized to soma. P-values were calculated using a Wilcoxon rank-sum test. ns (not significant) represents p > 0.05, * p < 0.05, ** p < 0.01, *** p < 0.001 and **** represents p < 0.0001.

Essentially all RP mRNAs share a highly conserved, pyrimidine-rich sequence motif in their 5*′* UTR called a 5*′* TOP (terminal <u>o</u>ligo<u>p</u>yrimidine) motif (Iadevaia et al., 2008; Yoshihama et al., 2002). We hypothesized that this shared motif could be responsible for the localization of RP mRNAs to the basal compartment of enterocytes. To test this, we selected two RP mRNAs, *RPL7* and *RPS28*, that were basally localized in the Halo-seq data (**Figure S3B**). We placed their entire 5*′* UTRs in front of the open reading frame (ORF) of Firefly luciferase to create two reporter transcripts (**Figure 3B**). The 5*′* UTRs of *RPL7* and *RPS28* are quite short (21 and 35 nucleotides, respectively), such that the entire 5*′* UTR is essentially a 5*′* TOP motif. As negative controls, we created reporter RNAs containing mutated versions of these 5*′* TOP motifs such that every pyrimidine was replaced with a purine (**Figure 3B**). We determined the localization of these reporter RNAs in C2bbe1 intestinal epithelial monolayers using smFISH probes designed to target the Firefly luciferase ORF.

On its own, the Firefly luciferase transcript is an exogenous, non-localized RNA. We found that addition of the 5*′* UTRs of both *RPL7* and *RPS28* were sufficient to drive the Firefly luciferase transcript to the basal compartment. Mutated 5*′* TOP motifs were not able to drive basal localization (**Figure 3C**). We then quantified the localization of each 5*′* TOP reporter’s transcripts and found that reporter transcripts containing 5*′* TOP motifs were significantly localized to the basal pole of the cells (p < 0.01, Wilcoxon rank sum test) while those containing mutant 5*′* TOP motifs were not (**Figure 3D**). To our knowledge, this is the first identified RNA element that is sufficient to drive RNAs to the basal pole of epithelial cells.

To more broadly assess the ability of the 5*′* TOP motif to regulate RNA localization in intestinal epithelial cells, we investigated the localization of our 5*′* TOP reporter constructs in a different human intestinal enterocyte cell line, HCA-7 cells. HCA-7 cells create monolayers that are more colon-like than the enterocyte-like C2bbe1 monolayers (Kirkland, 1985). Reporter constructs were integrated into HCA-7 lox cells via cre-*lox* recombination. We found that 5*′* TOP motifs from RPL7 and RPS28 were both sufficient for basal RNA localization in HCA-7 monolayers while mutated 5*′* TOP motifs were not (**Figure S3C**).

To test the ability of 5*′* TOP motifs to regulate RNA localization in epithelial cells generally, we integrated our human 5*′* TOP reporter constructs into MDCK cells through *cre*-mediated recombination. MDCK cells are epithelial cells derived from canine kidneys and have been used to study polarity across the apicobasal axis (Balcarova-Ständer et al., 1984). As before, we found that 5*′* TOP-containing reporter RNAs were strongly basally localized while mutant 5*′* TOP motifs were not able to drive basal RNA localization (**Figure S3D**). 5*′* TOP motifs are therefore able to promote basal RNA localization in a variety of epithelial cell types across multiple mammalian species. Furthermore, elements derived from human transcripts are able to regulate RNA localization in canine-derived cells.

### 5′ TOP motifs regulate RNA localization in morphologically dissimilar cell types

RP mRNAs have been found to be localized to cellular projections made by neurons or migrating cancer cells (Dermit et al., 2020; Shigeoka et al., 2019; Taliaferro et al., 2016) (**Figure S3A**). These observations were made by sequencing RNA fractions collected following mechanical fractionation using microporous membranes where small projections grow through pores restricting cell bodies on top of the membrane (**Figure 3E**). In these experiments RNA localization can be described as a localization ratio (LR) comparing the log2 abundance of a transcript in projection or neurite fraction to its abundance in the soma or cell body fraction (**Figure 3F**). To identify RNAs robustly localized into neurites, we combined published datasets from over 30 unique experiments where neuronal cells were subjected to mechanical fractionation. This dataset included neuronal samples spanning a range of neuronal cell types from cultured cell lines to primary cortical neurons (Goering et al., 2020; Taliaferro et al., 2016). For each neuronal sample, we calculated an LR value for each gene in each sample. These were then Z-normalized within a sample, and the gene’s LRz value was defined as the gene’s median normalized LR value across all samples. Genes with positive LRz values therefore have RNAs that are neurite-enriched while genes with negative LRz values have RNAs that are cell body-enriched. Using this dataset, we confirmed that RP mRNAs are efficiently localized to neurites (**Figure 3G**).

Although RP mRNAs are enriched in neuronal projections, the RNA sequences within the RP mRNAs that regulate this localization are unknown. We reasoned that because 5′ TOP motifs are highly conserved and had the ability to regulate RNA localization in epithelial cells that perhaps they may similarly be able to drive RNA transcripts to neurites. To test this, we integrated and expressed our human 5′ TOP reporter RNAs derived from RPL7 and RPS28 in CAD mouse neuronal cells. The endogenous mouse Rpl7 and Rps28 transcripts were found to be localized to neurites in the subcellular fractionation dataset (**Figure S3E**), and the 5′ TOP motifs of these mouse RNAs differ from their human orthologs by only a few nucleotides (**Figure S3F**). Using smFISH, we found that reporters with wildtype 5′ TOP motifs were efficiently localized to neurites while reporters that contained mutated 5′ TOP motifs or lacked 5′ TOP motifs were not (**Figure 3H**). We quantified the smFISH data by comparing the number of RNA puncta in the neurite to the number of puncta in the soma for each reporter. We found that 5′ TOP-containing reporters were significantly more enriched in neurites than reporters that contained mutated 5′ TOP motifs (p < 0.001, Wilcoxon rank-sum test) (**Figure 3I**). This effect is driven by a neurite-specific increase in transcript abundance and not due to any cell-wide RNA expression changes (**Figure S3G**). Human 5′ TOP motifs are therefore sufficient for neurite localization in mouse neuronal cells, suggesting that 5′ TOP-mediated RNA localization mechanisms are conserved across species. Further, given that 5′ TOP motifs regulate RNA localization in intestinal epithelial cells, neuronal cells, and migrating cancer cells (Dermit et al., 2020) (**Figure 3A, 3G, S3A**), these results demonstrate that 5′ TOP-mediated RNA localization mechanisms are conserved across cell types with vastly different morphologies.

### Neurite localization driven by 5′ TOP motifs is Larp1 dependent

We next sought to identify RBPs that are required for 5′ TOP-mediated RNA transport in neuronal cells. Multiple members of the highly conserved La superfamily of RBPs (also known as La Related Proteins or LARPs) regulate multiple aspects of RNA metabolism in a variety of cell types (Deragon, 2020; Dock-Bregeon et al., 2021). Two of these, Ssb and Larp1, have been previously found to bind 5′ TOP motifs (Dermit et al., 2020; Lahr et al., 2015; Maraia et al., 2017). Given this, we assayed the requirement of Larp family members for 5′ TOP-mediated RNA localization in CAD cells by performing a targeted RNAi screen and monitoring the localization of our 5′ TOP-containing reporters. Importantly, although Larp6 is required for endogenous RP mRNA transport in migrating cancer cell projections (Dermit et al., 2020), it is not expressed in CAD cells and therefore was not included in the screen.

We knocked down Larp family members Ssb, Larp1, Larp4, and Larp7 using siRNA transfection. Knockdown efficiencies assayed by qPCR (**Figure S4A**) and immunoblotting (**Figure 4A**) were between 80% and 95% for each Larp protein assayed. We hypothesized that if a Larp RBP was responsible for 5′ TOP-mediated RNA localization to neurites we would see a decrease in the neurite localization of 5′ TOP-containing reporters when expression of the Larp RBP was reduced. Following the depletion of Larp1, we observed that wildtype 5′ TOP-containing reporters were significantly less neurite-enriched, demonstrating that Larp1 is required for their efficient transport. This effect was 5′ TOP-specific as the localization of mutant 5′ TOP-containing reporters was unaffected (**Figure 4B-C**). Interestingly, Larp1 was the only assayed RBP that was required for 5′ TOP-mediated localization as knockdown of the other Larp family members did not consistently affect the localization of wildtype 5′ TOP reporters (**Figure S4B**). Larp1 therefore mediates 5′ TOP localization to neurites, and this function is not shared with other Larp family members.

**Figure 4.**
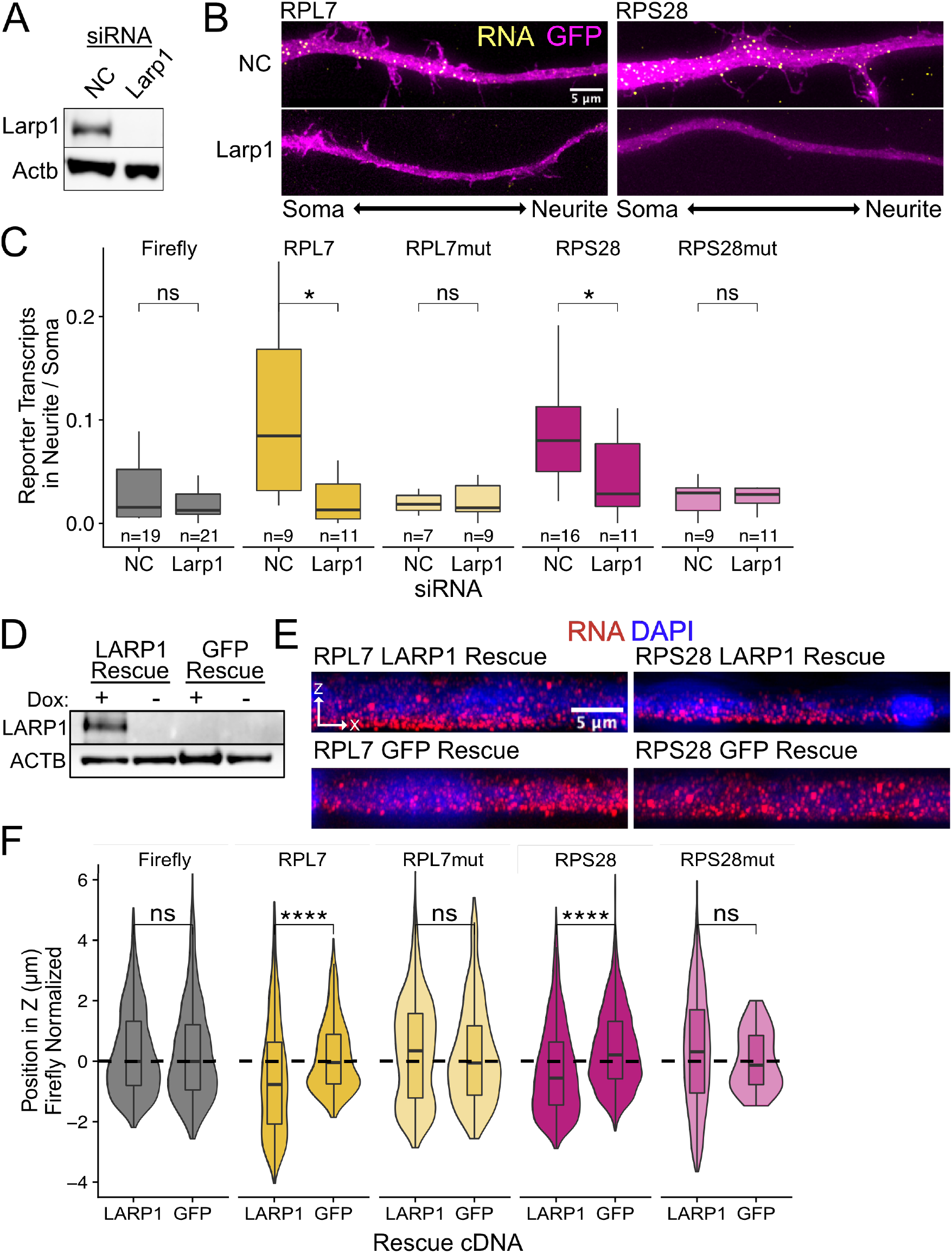
5′ TOP localization is mediated by LARP1 in enterocyte monolayers and neurons. (A) Immunoblot for Larp1 and Actb showing knockdown of Larp1 protein in neuronal cells when treated with Larp1 siRNAs as compared to a negative control (NC) siRNA. (B) Representative images of smFISH for each wildtype 5′ TOP reporter construct treated with Larp1 siRNAs or NC siRNAs. Images are a max projection through the XY axis of a single neurite positioned with the soma to the left. RNA signal shown in yellow while cell outlines marked by GFP signal shown as magenta. (C) Number of smFISH reporter transcript puncta in neurites normalized to soma. P-values were calculated using a Wilcoxon rank-sum test. (D) Immunoblot for LARP1 and ACTB showing knockout and doxycycline-induced rescue of LARP1 levels in C2bbe1 LARP1 knockout cells. (E) Representative images of smFISH for each wildtype 5′ TOP reporter construct. RNA signal shown in red while DAPI marks the nuclei in blue. Images are a max projection through the XZ axis of many cells. (F) smFISH puncta position in Z of the indicated Firefly luciferase reporters normalized to median untagged Firefly luciferase transcript position. P-values were calculated using a Wilcoxon rank-sum test. ns (not significant) represents p > 0.05, * p < 0.05, ** p < 0.01, *** p < 0.001 and **** represents p < 0.0001.

### Larp1 depletion reduces number and length of neurites

During our Larp depletion experiments, we observed that the loss of Larp1 resulted in CAD cells with defects in neurite outgrowth (**Figure S4C**). Specifically, the loss of Larp1 expression reduced both the number of cells producing neurites (**Figure S4D**) and shortened overall neurite length (**Figure S4E**). Larp1 was the only tested Larp protein for which this effect was observed. The depletion of other Larp proteins increased the number of cells producing neurites (**Figure S4D**), and in the case of Larp4, also increased overall neurite length (**Figure S4E**). Larp1 is therefore both required for transport of 5′ TOP-containing transcripts to neurites, and loss of Larp1 inhibits neurite development and neurite lengthening. However, because many Larp proteins, including Larp1, are implicated in a wide range of RNA regulatory modes, we cannot directly ascribe this neurite phenotype to the loss of RP mRNA localization.

### Basal localization of 5′ TOP motifs in epithelial cells is LARP1 dependent

Following our knockdown experiments in mouse neuronal cells, we suspected that the TOP-binding protein LARP1 may be responsible for the basal localization of 5′ TOP-containing RNAs in human epithelial monolayers. To test this, we used CRISPR/Cas9 to generate a LARP1 knockout C2bbe1 cell line.

Using two guide RNAs, we targeted exon 2 and exon 19, both of which are included in all major isoforms of LARP1, with the goal of deleting the sequence in between them (**Figure S4F**). Because C2bbe1 cells may not be diploid at the *LARP1 locus*, we expected to identify multiple Cas9-generated *LARP1* alleles in a single clonal line. We identified one clonal line in which no wild type *LARP1* was detectable. One or more alleles had lost the sequence in between exon 2 and exon 19 and the only remaining allele contained a 53 base pair deletion in exon 2 that results in a premature stop codon (**Figure S4F, G**). The *LARP1* RNA produced by this clone is therefore not expected to produce a full length protein product and should be targeted for nonsense mediated decay (NMD) (**Figure S4F**). Consistent with this, the level of *LARP1* RNA produced in this clone was less than 3% of that produced in wildtype cells (**Figure S4H**).

To reintroduce expression of LARP1 in the selected LARP1 knockout clone, a full length LARP1 rescue cDNA under a doxycycline-inducible promoter was integrated via *cre-*mediated recombination. This LARP1 KO-rescue line expresses LARP1 protein only in the presence of doxycycline. As a control, we also created a line in which the knockout was rescued with doxycycline-inducible GFP (**Figure 4D**). Rescuing with LARP1 cDNA restores any LARP1 localization function while rescuing with GFP cDNA maintains LARP1 knockout status.

By expressing our 5′ TOP-containing reporter RNAs in the LARP1 knockout and rescue lines, we found that 5′ TOP-containing reporters were basally localized only when LARP1 was present, indicating that LARP1 is required for their localization (**Figure 4E,F**). Altogether, we found that LARP1 medicates the basal localization of 5′ TOP-containing RNAs in epithelial monolayers and the neurite localization of the same RNAs in neuronal cells.

### TOP motifs must be at the 5′ end of transcripts in order to mediate RNA localization

To better understand how 5′ TOP motifs were regulating the localization of our reporter transcripts, we asked if the position of the motif in the transcript changed its ability to regulate RNA localization. Efficient binding of LARP1 to 5′ TOP motifs requires close proximity to the 5′ 7-methylguanosine cap (Al-Ashtal et al., 2021; Lahr et al., 2017; Philippe et al., 2018). To test the requirement of TOP proximity to the 5′ 7-methylguanosine cap for RNA localization regulation, we placed the RPL7 and RPS28 TOP motifs directly after the Firefly luciferase ORF, thus creating 3′ TOP reporter constructs (**Figure 5A**). We reasoned that if the 5′ location of the TOP motif was important, the 3′ TOP reporters would display a reduced ability to regulate RNA localization. In C2bbe1 monolayers, we found that 3′ TOP reporters were significantly less basally localized than their 5′ TOP counterparts, indicating that the position of the TOP motif within transcripts is important for their RNA localization regulatory activity (**Figure 5B**). We observed similar results in mouse neuronal cells, finding that 3′ TOP reporters were significantly less efficient at driving RNAs to neurites than their 5′ TOP counterparts (**Figure 5C**). Therefore, in order to efficiently regulate RNA localization, TOP motifs must be at the 5′ end of transcripts. This is consistent with LARP1 binding both TOP motifs and the 5′ 7-methylguanosine cap (Al-Ashtal et al., 2021; Lahr et al., 2017; Philippe et al., 2018).

**Figure 5.**
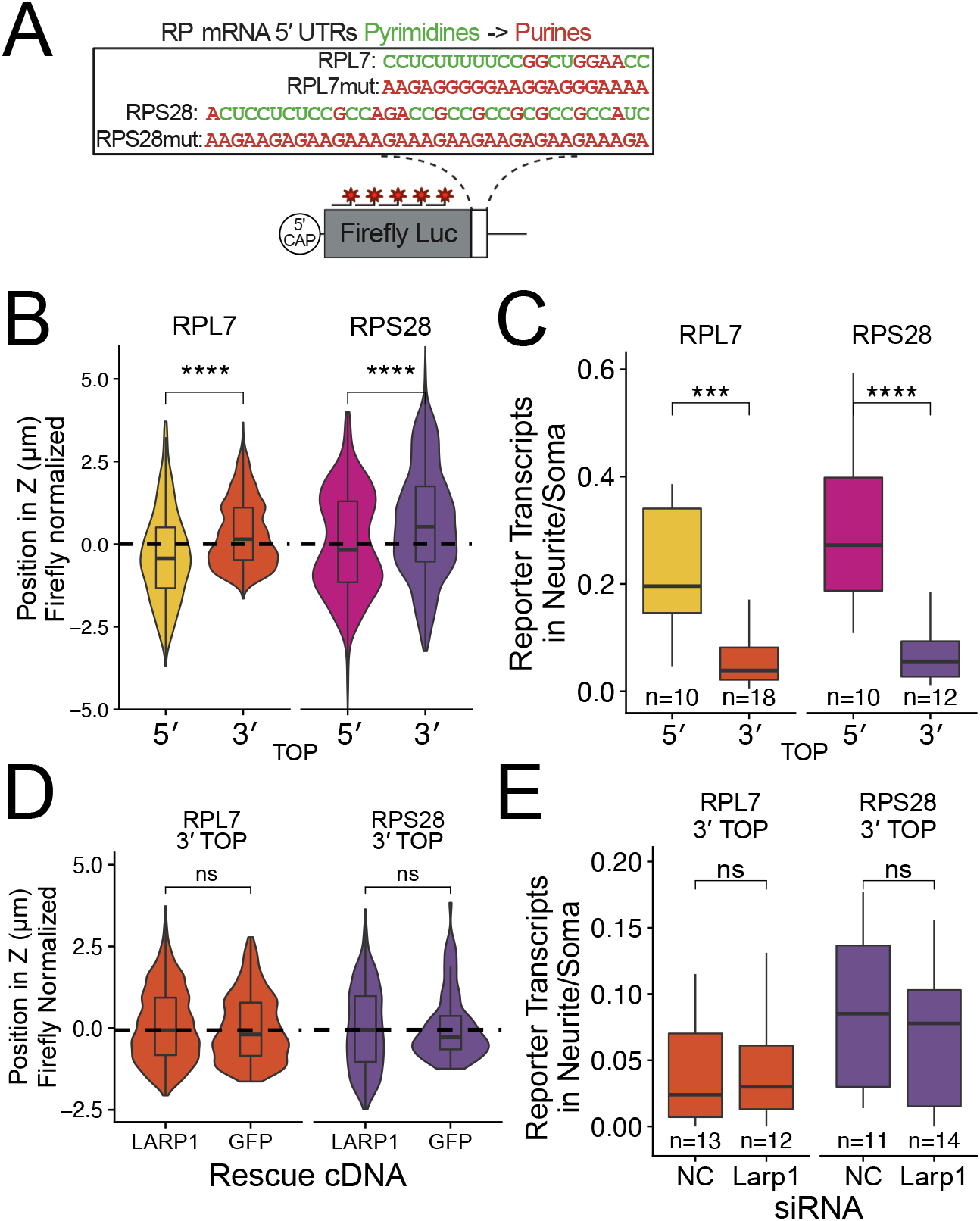
5′ TOP motif position in transcript is required for efficient transcript localization. (A) Schematic of ribosomal protein 3′ TOP reporter constructs. Mutant versions were created by swapping each pyrimidine for a purine. TOP motif sequences are identical to 5′ TOP reporter constructs. (B) 3′ TOP reporter transcript localization in C2bbe1 monolayers. smFISH puncta position of the indicated Firefly luciferase reporters in Z normalized to median untagged Firefly transcript position. P-values were calculated using a Wilcoxon rank-sum test. (C) 3′ TOP construct localization in neurons. Number of reporter RNA smFISH puncta in neurites normalized to soma. P-values were calculated using a Wilcoxon rank-sum test. (D) 3′ TOP construct localization in LARP1 knockout and rescued C2bbe1 monolayers. smFISH puncta position of the indicated Firefly luciferase reporters in Z normalized to median untagged Firefly luciferase transcript position. P-values calculated using a Wilcoxon rank-sum test. (E) 3′ TOP construct localization in neurons depleted of Larp1 by siRNA. Number of reporter RNA smFISH puncta in neurites normalized to soma. P-values calculated using a Wilcoxon rank-sum test. ns (not significant) represents p > 0.05, * p < 0.05, ** p < 0.01, *** p < 0.001 and **** represents p < 0.0001.

Interestingly, the mutant purine-rich 3′ TOP reporter transcripts were strongly basally localized in epithelial cells (**Figure S5A**). We found similar results in neuronal cells where the RPL7 3′ TOP reporter RNA was neurite-localized (**Figure S5B**). Computational prediction of RNA secondary structure revealed that the mutant RPL7 TOP motif is likely folded into a G-quadruplex conformation (Lorenz et al., 2011) (**Figure S5C**). G-quadruplex structures within 3′ UTRs have been previously found to target RNAs to neurites (Goering et al., 2020; Subramanian et al., 2011) and may therefore be similarly regulating RNA localization in epithelial cells.

To ask if LARP1 regulates the localization of 3′ TOP-containing transcripts, we assayed 3′ TOP localization in LARP1 and GFP rescued LARP1 KO C2bbe1 monolayers using smFISH. We found that the loss of LARP1 did not affect the localization of 3′ TOP-containing transcripts (**Figure 5D, S5D**). Similarly, in mouse neuronal cells, siRNA-mediated depletion of Larp1 did not affect the neurite localization of 3′ TOP-containing transcripts (**Figure 5E, S5E**). These results further strengthen the argument that LARP-1 mediated RNA localization requires a TOP motif very near the 5′ terminus of the transcript.

### An RNA element within the 3′ UTR of Net1 is necessary and sufficient for RNA localization in neuronal and epithelial cells

We had identified an RNA element, 5′ TOP motifs, that regulated RNA localization in both epithelial and neuronal cells. We wondered whether other RNA elements may be similarly able to regulate RNA localization in both cell types. We therefore set out to find other examples of RNAs that were localized in both cell types as well as the sequences that drive their localization.

*Net1* RNA encodes a guanine exchange factor and is strongly localized to neurites (Arora et al., 2021) and the basal pole of intestinal epithelial cells (Moor et al., 2017). The element responsible for the localization of mouse *Net1* RNA to neurites has recently been identified as a GA-rich region in the transcript’s 3′ UTR (Arora et al., 2021). The 3′ UTR of the human *NET1* transcript also contains a highly GA-rich region (Arora et al., 2021).

To ask if the same RNA sequences in *Net1* RNA that drive RNA localization in mouse neuronal cells are also active in human epithelial cells, we created reporter transcripts that contained 3′ UTR sequences derived from mouse *Net1*. These reporter constructs either contained the full length *Net1* 3′ UTR, just the identified GA-rich localization element (LE) or the entire *Net1* 3′ UTR lacking the LE (ΔLE) (**Figure 6A**). As before, the reporter constructs were integrated into the genomes of C2bbe1 cells, inducibly expressed with doxycycline, and the localization of the resulting transcripts was visualized using smFISH. We found that the full length *Net1* 3′ UTR was sufficient to strongly drive RNAs to the basal pole of epithelial cells. The LE displayed significant yet weaker RNA localization activity, and the 3′ UTR lacking the localization element (ΔLE) displayed no basal localization activity (**Figure 6B, D**). These results indicate that the GA-rich LE within the mouse *Net1* 3′ UTR is both necessary and sufficient to drive RNA to the basal pole of human enterocytes.

**Figure 6.**
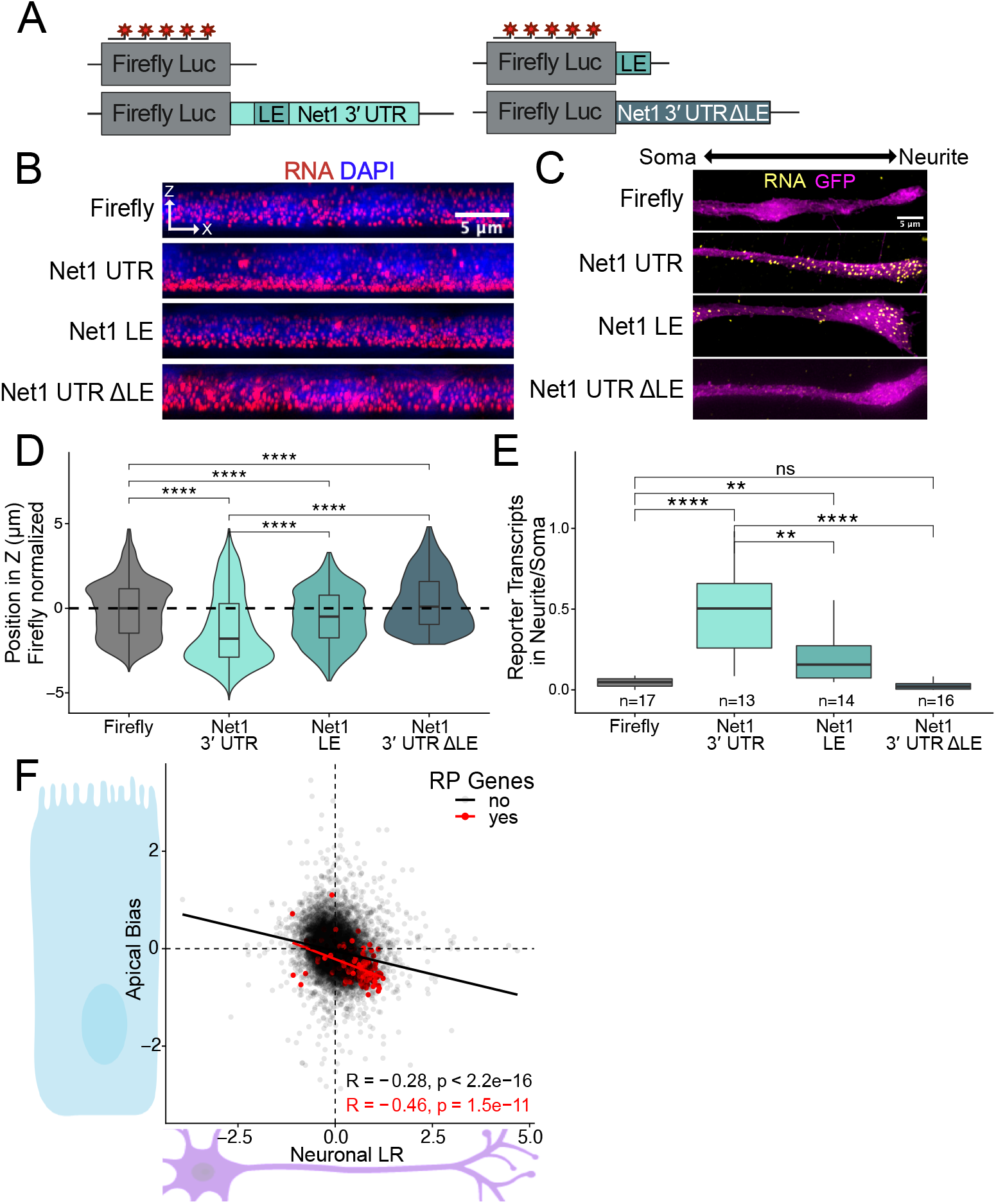
RNA localization mechanisms beyond 5′ TOP motifs are conserved across species and cell morphology. (A) Schematic of Net1 reporter constructs used to investigate necessity and sufficiency of Net1’s localization element (LE). (B) Representative images of smFISH puncta for each Net1 reporter construct in C2bbe1 monolayers. RNA signal shown in red while DAPI marks the nuclei in blue. Images are a max projection through the XZ axis of many cells. (C) Representative images of smFISH puncta for Net1 constructs in CAD cells. Images are a max projection through the XY axis of a single neurite positioned with the soma to the left. RNA signal shown in yellow while cell outlines marked by GFP signal shown as magenta. (D) smFISH puncta position in Z of the indicated reporter RNAs normalized to median untagged Firefly luciferase transcript position. P-values were calculated using a Wilcoxon rank-sum test. (E) Number of reporter RNA smFISH puncta in neurites normalized to soma. P-values were calculated using a Wilcoxon rank-sum test. (F) Direct comparison of the Halo-seq-derived Apical Bias values in human C2bbe1 monolayers and mechanical fractionation-derived neuronal Localization Ratio (LR) values in neuronal cells for all genes. Ribosomal protein mRNAs are highlighted in red. ns (not significant) represents p > 0.05, * p < 0.05, ** p < 0.01, *** p < 0.001 and **** represents p < 0.0001.

We then assayed the same reporter constructs in mouse neuronal CAD cells using smFISH. As in the human epithelial cells, we found that the full length *Net1* 3′ UTR strongly drove transcripts to neurites, the LE had significant but reduced activity, and the ΔLE 3′ UTR displayed no activity (**Figure 6C, E**). These results were due to a neurite-specific accumulation of reporter transcripts and not an overall difference in transcript expression (**Figure S6A**). The GA-rich sequence element within the 3′ UTR of mouse *Net1* therefore represents another sequence that regulates RNA localization across cell types, species and cellular morphologies.

### Enterocytes and neurons have similar localized transcriptomes

If similar mechanisms were transporting RNAs to the basal pole of epithelial cells and the neurites of neuronal cells, we reasoned that these two subcellular compartments should be enriched for similar RNAs. To test this, we compared LRz and AB values from the neuronal fractionation and epithelial Halo-seq datasets, respectively, transcriptome-wide (**Figure 6F**). Somewhat surprisingly, we found a reasonably strong and highly significant correlation between these values (R = -0.28, p < 2.2e-16). The negative sign of this correlation indicates that neurites and the basal pole of epithelial cells are enriched for similar transcripts. RP mRNAs, highlighted in red, exemplify this relationship (R = -0.46, p = 1.5e-11). These correlations are even more striking when considering the fact that the LRz and AB values are derived from experiments that differ greatly in their approaches (mechanical subcellular fractionation vs. proximity labeling), species (mouse vs. human), and cell types (neurons vs. intestinal epithelial cells). These very different cellular compartments are therefore spatial equivalents, at least in terms of RNA localization, and their RNA contents are likely governed by the same regulatory mechanisms.

The genes that fall in the fourth quadrant of Figure 6F are localized basally in enterocytes and to the neurites of neurons. To interrogate the identity of these genes we used GO enrichment analysis (Alexa et al., 2010). The most enriched GO terms were “ribosome” and “mitochondrion” (**Figure S6B**). We have extensively detailed that RP mRNAs are enriched both in neurites and the basal compartment of epithelial cells (**Figure 3A, 3G, S3A**). We have similarly found that mitochondria are enriched toward the basal pole of epithelial cells (**Figure S2H**), and mitochondria have previously been reported to be found in neuronal projections (Mandal and Drerup, 2019). Given that many nuclear-encoded RNAs that encode mitochondrial proteins are translated on the surface of the mitochondria, it may be expected that these RNAs are also neurite- and basal-enriched. Consistent with this idea, we found that RNAs that encode components of the electron transport chain (ETC) were both neurite- and basal-enriched according to our sequencing data sets (**Figure S6C**). This argues that similarities in RNA localization between these two cell types can be driven by similar organelle localization.

### RNA localization to neurites and the basal pole of epithelial cells is dependent on kinesin activity

Despite drastically different morphologies, intestinal epithelial cells and neurons are organized using similar underlying microtubule structures. These polar molecules are organized in such a way that their plus ends are oriented basally in intestinal epithelial cells and out to neurite tips in neurons (Müsch, 2004; Sugioka and Sawa, 2012; Yogev et al., 2016) (**Figure 7A**). Because of the similar microtubule architecture in both cell types and the fact that RNAs are known to be trafficked along microtubules (Cross et al., 2021; Gagnon and Mowry, 2011), we hypothesized that RNAs were being trafficked to neurites and the basal pole of epithelial cells through the action of kinesin motor proteins. To test this hypothesis, we monitored the localization of our reporter RNAs following treatment of cells with the kinesin inhibitor kinesore (Randall et al., 2017).

**Figure 7.**
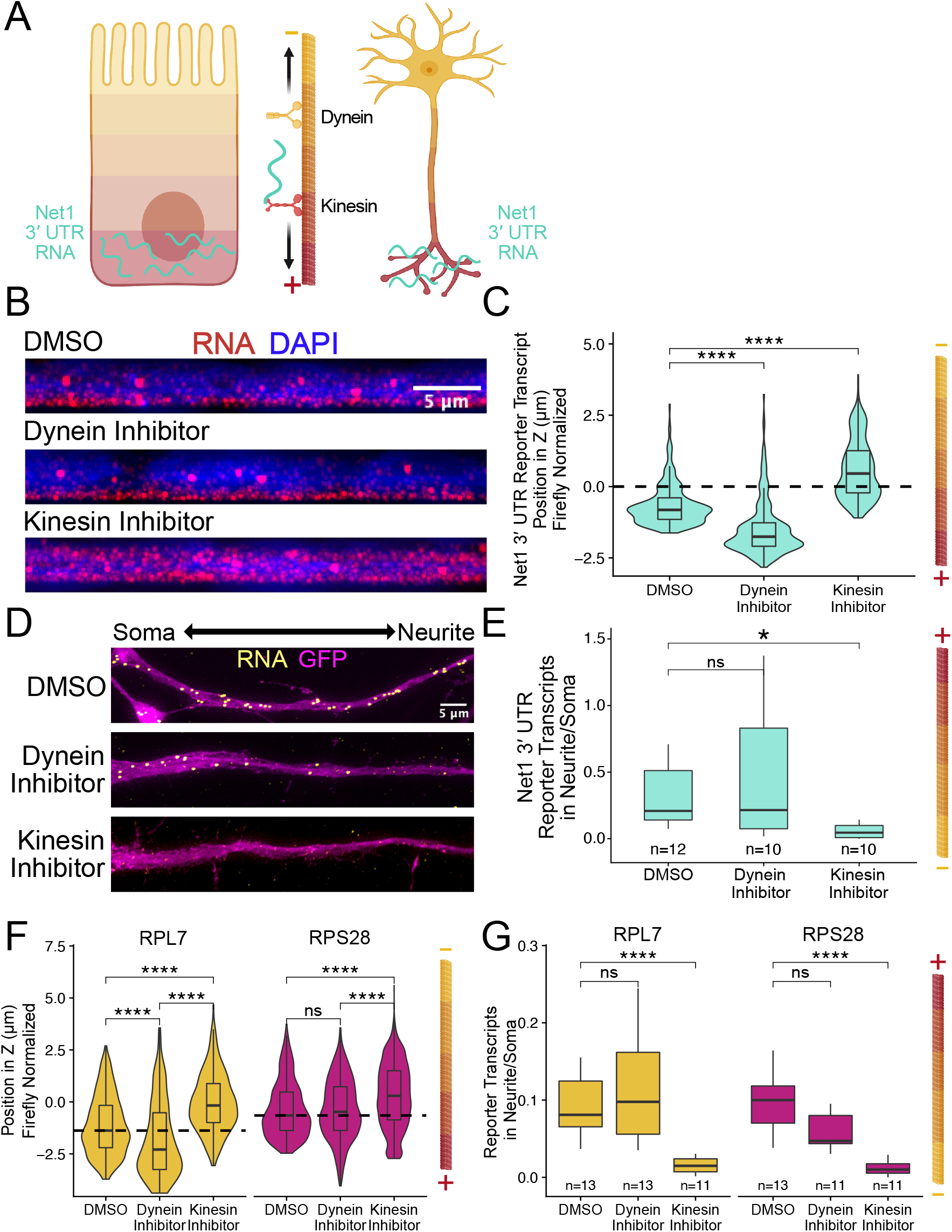
RNA localization to neurites and the basal pole of epithelial cells is dependent on kinesin. (A) Schematic of microtubule organization in both enterocytes and neurons where the + ends are oriented basally and out to neurites. Net1 3′ UTR Reporter constructs are teal, hypothesized to reach their destination via kinesin transport. (B) Representative images of Net1 3’UTR reporter transcript localization with each inhibitor treatment or DMSO control. RNA signal shown in red while DAPI marks the nuclei in blue. Images are a max projection through the XZ axis of many cells. (C) smFISH puncta position in Z of Firefly luciferase-Net1 3′ UTR constructs normalized to median untagged Firefly luciferase transcript position in Z. P-values were calculated using a Wilcoxon rank-sum test. (D) Representative images of smFISH for Net1 3’UTR transcripts in CAD cells. Images are a max projection through the XY axis of a single neurite positioned with the soma to the left. RNA signal shown in yellow while cell outlines marked by GFP signal shown as magenta. Number of reporter RNA smFISH puncta in neurites normalized to soma. P-values were calculated using a Wilcoxon rank-sum test. smFISH puncta position in Z of RPL7 and RPS28 5′ TOP constructs normalized to median untagged Firefly luciferase transcript position in Z. P-values were calculated using a Wilcoxon rank-sum test. (G) Number of reporter RNA smFISH puncta in neurites normalized to soma. P-values were calculated using a Wilcoxon rank-sum test. ns (not significant) represents p > 0.05, * p < 0.05, ** p < 0.01, *** p < 0.001 and **** represents p < 0.0001.

Cells expressing the full length *Net1* 3′ UTR reporter construct were treated for four hours with kinesore (Randall et al., 2017), a kinesin-1 inhibitor, or ciliobrevin A, a dynein-1 inhibitor. In C2bbe1 monolayers, we found that kinesin inhibition dramatically reduced the basal localization of reporter RNAs containing the *Net1* 3′ UTR (**Figure 7B,C)**, consistent with the hypothesis that these RNAs are trafficked along microtubules in epithelial cells. Interestingly, we found that inhibition of dynein strengthens basal localization of Net1 3′ UTR transcripts. Kinesin and dynein activity are often closely linked, pulling the same cargoes in different directions. When the activity of one motor is inhibited, it can therefore increase the apparent activity of the other (Soppina et al., 2009). In neuronal cells, kinesin inhibition again significantly reduced the neurite localization of the *Net1* 3′ UTR reporter RNAs, again suggesting they are transported along microtubules (**Figure 7D,E**). We did not observe any changes in control Firefly luciferase reporter RNA localization in response to drug treatments (**Figure S7A,B**). Additionally, as before, changes in RNA localization were driven by changes in RNA content of the neurite and not any gross changes in expression (**Figure S7C**).

We found very similar results with our 5′ TOP reporter transcripts when inhibiting motor proteins. In C2bbe1 monolayers, kinesin inhibition removed the basal localization of both RPL7 and RPS28 5′ TOP reporter constructs (**Figure 7F**). Kinesin mediated transport along microtubules was also required for the neurite localization of 5′ TOP containing transcripts (**Figure 7G**). Once again, changes in RNA abundance were neurite-specific and were not observed cell-wide (**Figure S7D**).

Given that the plus ends of microtubules are associated with both neurites and the basal pole of epithelial cells and that kinesin inhibition reduced RNA localization to both compartments, we conclude that the RNA contents of both locations are determined at least in part through microtubule plus-end directed trafficking mechanisms.

## DISCUSSION

For the vast majority of RNAs that display a particular subcellular localization, the RNA elements that regulate this localization are unknown (Engel et al., 2020). For those that are known, their activity has generally only been studied in a single cell type (e.g. neurons or *Drosophila* embryos). Because the subcellular location of an RNA is often intricately linked with the morphology of the cell, it has been difficult to predict how RNA localization elements that are active in one cell type would behave in another. In this study we have demonstrated that two RNA localization elements, 5′ TOP motifs and a GA-rich element in the 3′ UTR of the mouse *Net1* RNA, drive RNAs to the projections of mouse neuronal cells and the basal pole of human epithelial cells. Further, transcriptome-wide analysis of these subcellular compartments revealed that their RNA contents are similar, suggesting that the same mechanisms are populating their localized transcriptomes.

RP mRNAs have been previously identified as enriched in neurites (Taliaferro et al., 2016) and the basal pole of epithelial cells (Moor et al., 2017). However, the mechanisms that direct RP mRNAs to those locations have been unknown. We found that 5′ TOP motifs regulate the trafficking of RNAs to both of those locations and do so in a manner that requires the RBP LARP1. 5′ TOP motifs were also previously found to regulate RNA localization to the protrusions of migrating cells (Dermit et al., 2020). However, in these cells, RP mRNA localization required another La-family RBP, LARP6. Larp6 was not expressed in our mouse neuronal model, and LARP6 was only lowly expressed in our human epithelial model. Thus, it is possible that LARP1 and LARP6 have redundancy in RNA localization activities and can complement each other in this regard if expressed in the same cell.

Interactions between LARP1 and 5′ TOP motifs have been well-studied (Al-Ashtal et al., 2021; Lahr et al., 2015, 2017). Generally, these interactions have been characterized in the context of LARP1’s ability to regulate the translational status of 5′ TOP-containing mRNAs (Lahr et al., 2017). We report here a new function for LARP1, the regulation of RNA localization. Importantly, it has been demonstrated that LARP1 contains at least three RNA binding regions, a DM15 region that recognizes the 5′ mRNA cap, an RNA recognition motif and the La motif that recognizes the pyrimidine-rich TOP motif. RNA binding is tightest when both of these binding modes (i.e. to the 5′ cap and the TOP motif) are engaged (Lahr et al., 2017). We found that TOP motifs, when placed in the 3′ UTR of a transcript, were unable to regulate the localization of the transcript, both in neuronal and epithelial cells. Given that we identified LARP1 as critical for this process and that LARP1 requires the TOP motif to be adjacent to the 5′ cap for efficient RNA binding, we propose that LARP1 is not able to efficiently recognize the 3′ TOP motifs, resulting in their reduced transcript localization.

Intriguingly, the depletion of Larp1 in mouse neuronal cells and the knockout of LARP1 in human epithelial cells led to noticeable phenotypes. In neuronal cells, Larp1 depletion led to cells producing fewer and shorter neurites. LARP1 knockout epithelial cells had a noticeable growth defect and had considerably longer doubling times. It is tempting to ascribe at least parts to these phenotypes to the mislocalization of RP mRNAs, especially since RP mRNAs are some of the most highly expressed mRNAs in the cell. However, given that LARP1 participates in multiple facets of post-transcriptional gene regulation (Berman et al., 2021), we cannot directly make this conclusion.

The conserved localization of RP mRNAs as a class across multiple species and cell types suggests a repeated need for the spatial control of ribosomal protein synthesis. The translation of RP mRNAs could greatly impact the translational output of an entire cell as more ribosomal proteins allows for a greater number of functional ribosomes (Fonseca et al., 2018). Still, given that ribosome assembly canonically occurs in the nucleolus, it is puzzling as to why RNAs encoding ribosomal subunits would often be kept so far away from the nucleus. One idea is that locally synthesized ribosomal proteins can directly join and remodel, or perhaps repair, nearby ribosomes, promoting further local control of translation (Shigeoka et al., 2019).

We also found that a GA-rich region in the 3′ UTR of the mouse *Net1* RNA is necessary and sufficient to drive RNAs both to the neurites of mouse neuronal cells and the basal pole of human epithelial cells. Interestingly, a GA-rich region in the 3′ UTR of human *NET1* RNA was also identified as necessary for the localization of the transcript to the leading edges of migrating cells (Chrisafis et al., 2020)

Overall, these findings point to the existence of an RNA localization “code” that operates in multiple cell types and is conserved across species. In the mechanisms illuminated in this study, we propose that the RNA localization codes we identified are not “neurite localization” signals or “basal localization” signals *per se*. Rather, they are signals that specify that an RNA should be trafficked to the plus end of microtubules, wherever that may be in a given cell type. This opens up the possibility of predicting the localization of an RNA in a given cell type if its localization in another cell type is known, greatly increasing our understanding of mechanisms underlying RNA trafficking.

## ACKNOWLEDGEMENTS

We thank members of the Taliaferro lab for helpful discussions regarding experiments and analyses. We further thank Rytis Prekeris and Peter Dempsey for helpful discussions and advice regarding epithelial cell biology. Portions of the figures in this manuscript were created using Biorender.

This work was funded by the National Institutes of Health (R35-GM133885 to JMT), the W.M. Keck Foundation (JMT) and the RNA Bioscience Initiative at the University of Colorado Anschutz Medical Campus (JMT, RG).

This work was further supported by a Predoctoral Training Grant in Molecular Biology NIH-T32-GM008730 to RG).

## METHODS

### Cell maintenance, differentiation

C2bbe1 cells (CRL-2102) were maintained in DMEM with 110 mg/L Sodium Pyruvate (Thermo 11995065) supplemented with 10% FBS (Atlas Biologicals F-0500-D), penicillin-streptomycin (Thermo 15140122) and 10 ug/mL human transferrin (Sigma-Aldrich T8158). HCA-7 cells were maintained in a 1:1 mix of DMEM and F12 (Thermo 11320033) supplemented with 10% FBS (Atlas Biologicals F-0500-D) and penicillin-streptomycin (Thermo 15140122). MDCK cells were maintained in DMEM (Thermo 11965092) supplemented with 10% Equafetal (Atlas Biologicals EF-0500-A) and penicillin-streptomycin (Thermo 15140122).

C2bbe1 LoxP cells were maintained in 20 µg/mL blasticidin (A.G. Scientific B-1247-SOL) while HCA-7 and MDCK lox cells were maintained in 10 µg/mL blasticidin. Following *cre-*mediated recombination with a puromycin resistance plasmid, all reporter-expressing lines were maintained in 5 µg/mL puromycin (Cayman Chemical Company 13884).

C2bbe1, HCA-7 and MDCK cells were allowed to differentiate for 7 days after plating at 100% confluency in the presence of 2 µg/mL doxycycline (Fisher Scientific AAJ6042203) when inducing transgene and/or reporter transcript expression. Differentiation can occur in dishes, plates or on 0.4 µm transwell inserts (Corning 353090). Media was changed every 3-4 days.

CAD cells were maintained in a 1:1 mix of DMEM and F12 (Thermo 11320033) supplemented with 10% Equafetal (Atlas Biologicals EF-0500-A) and penicillin-streptomycin (Thermo 15140122). CAD lox cells were received as a gift from Eugene Makeyev (Khandelia et al., 2011) and maintained in 5 µg/mL blasticidin (A.G. Scientific B-1247-SOL).

CAD cells were grown in full growth media and induced with 2 µg/mL doxycycline for 48 hours when inducing expression of transgenic constructs. Following doxycycline induction, media was replaced with serum free media for 48 hours which induces neurite outgrowth. Doxycycline induction was continued through the differentiation period when inducing construct expression. Reporter-expressing CAD lines were maintained in 5 µg/mL puromycin (Cayman Chemical Company 13884).

### CRISPR/Cas9 modifications

A loxP-flanked blasticidin resistance cassette was integrated into the AAVS1 safe harbor of C2bbe1 and HCA-7 cells via CRISPR/Cas9. RNPs from Synthego with AAVS1 targeting sgRNA (UAGUGGCCCCACUGUGGGGU) were electroporated with the Neon electroporation system (C2bbe1:1600V 10ms and 3 pulses, HCA-7: 1100V 20ms 2 pulses). A donor plasmid containing the loxP cassette with homology to the AAVS1 safe harbor was co-electroporated with the RNPs. Following electroporation, cells were incubated at 37°C for 48 hours before selection with blasticidin (20 µg/mL for C2bbe1, 10 µg/mL for HCA-7) until untransfected negative control cells had died and the newly selected loxP line exhibited normal growth patterns.

MDCK loxP cells were created through random integration using the same electroporation (1600V 10ms 3 pulses) strategy and donor plasmid for human lines. CRISPR/Cas9 RNPs were omitted from this electropporation. Selection of blasticidin resistant clones contained loxP cassettes in undetermined locations.

For the LARP1 knockout line, C2bbe1 loxP cell lines were treated with two sgRNAs from Synthego that targeted exon 2 (GCUGUUCCUAAACAGCGCAA) and exon 19 (GCUGUUCCUAAACAGCGCAA) of LARP1. CRISPR Cas9 RNPs and a GFP plasmid were co-electroporated using the Neon electroporation system (1600V 10ms and 3 pulses) and incubated at 37°C for 48 hours. GFP positive cells were sorted to single cells using a flow cytometer (MoFlo XDP100) and allowed to expand for 2-4 weeks. Clones were screened for gene deletion by PCR as well as LARP1 mRNA and protein depletion determined by qPCR and western blot analysis with a monoclonal LARP1 antibody (Santa Cruz sc-515873). Lesions at the locus of cutting were identified by PCR using genomic DNA to amplify the locus followed by Sanger sequencing and Synthego ICE analysis.

### Plasmid construction

All plasmids are derivatives of pRD-RIPE (Khandelia et al., 2011). All Halo-fusion proteins were cloned into the location of the GFP ORF by excising GFP with AgeI and BstXI and then PCR amplifying the cDNA of the Halo fusion protein of interest with overhanging primer handles that provide 20 bp of homology to the plasmid backbone. Some Halo fusion cDNAs were synthesized by Twist Bioscience. PODXL (Horizon, MHS6278-202858197), ATP1A1 (Horizon, MHS6278-202759485), and LARP1 (Horizon, MHS6278-202827213) cDNAs were obtained from Horizon Discovery. Infusion cloning (TakaraBio) was used to ligate PCR amplified inserts into plasmid backbones. Constructs were confirmed by restriction digest assays and DNA sequencing.

A variant of pRD-RIPE with a bi-directional promoter driving both Firefly and Renilla luciferases was inserted into pRD-RIPE creating pRD-RIPE-BiTET-FR. Reporter constructs were created by adding additional sequence to cut sites in the 5′ (Asc1) or 3′ (PmeI) end of the Firefly luciferase ORF. Because TOP motifs are short, DNA inserts were created by annealing two oligos and cloning via sticky end cloning with T4 DNA Ligase (NEB M0202S) into the linearized pRD-RIPE-BiTET-FR.

### Cre recombinase-mediated cassette switching

All loxP-containing cell lines followed the same protocol for cassette switching. Cells were plated in 12-well plates at 1.0-1.5 × 105 cells per well in full growth media 12-18 hours before transfection. Cells were then co-transfected with 500 ng donor cassette plasmid with 1% (wt/wt) of a Cre-encoding plasmid (pBT140, addgene #27493). Plasmids were transfected using Lipofectamine LTX reagent (Thermo Fisher Scientific 5338030) and Opti-MEM (Thermo Fisher Scientific 31985070) following the manufacturer’s protocol. Following 24 hours with transfection reagents, the medium was changed and the incubation continued for another 24 hours before addition of 1-5 µg/mL puromycin (VWR 97064-280). Incubation with puromycin continued for several days until untransfected control wells were depleted of cells. The puro-resistant colonies were expanded in a full growth medium supplemented with 5 µg/mL puromycin.

### Halo SDS-PAGE gel stain

Halo-expressing cells were grown in the presence or absence of 2 µg/mL doxycycline for 48 hours before lysate collection. Following a PBS wash, cells were incubated with 25 nM Halo ligand Janelia Fluor 646 (Promega GA1120) for 10 minutes. Cells were washed 3 times with PBS. Lysates from 1.0-1.5 × 106 cells were scraped into 50-75 µL RIPA lysis buffer (50 mM Tris-HCl, pH 7.0, 150 mM NaCl, 0.1% (w/v) SDS, 0.5% sodium deoxycholate, 1.0% Triton X, 5 mM EDTA). Lysates were chilled on ice for 15 minutes before sonication for 30 seconds. 15 µL of lysates were mixed with sample buffer (Invitrogen NP0008) before being incubated at 100°C for 5-10 minutes. Samples were loaded into 4-12% Bis-Tris 1mm SDS protein gel (Thermo Fisher Scientific NP0323BOX) with protein ladder (Gel Company FPL-008). Gels were imaged with a Sapphire molecular imager (Azure Biosystems) set to collect fluorescent Cy5 signal. Following fluorescent imaging the gel was stained with 1% Coomassie (VWR 95043-420) for 20 minutes, de-stained (40% Methanol, 10% Acetic Acid) 1-16 hours, and imaged with the visible setting.

### Halo Imaging in fixed cells

Halo fusion proteins were visualized in cells by staining with fluorescent Halo ligand (Promega GA1110). Cells were seeded on polyD lysine coated coverslips (neuVitro) in full growth media supplemented with 2 µg/mL doxycycline to induce Halo transgene expression for 48 hours. Following a PBS wash, cells were incubated with Halo ligand Janelia Fluor 549 (Promega) at 25 nM for 30 minutes at room temperature. Following PBS wash, cells were fixed with 3.7% formaldehyde for 20 minutes at room temperature. Cell nuclei were stained with 100 ng/mL DAPI (Sigma-Aldrich D9542) for 15 minutes, and then cells were washed again with PBS. Coverslips were mounted with fluoromount G (SouthernBiotech 0100-01) and imaged at 60X magnification with a Deltavision Elite widefield fluorescence microscope (GE).

### Halo-seq

Halo-seq was performed as previously described (Engel et al., 2021). Details regarding each step of Halo-seq are provided below.

#### In-cell labeling

Halo transgene-expressing C2bbe1 cells were seeded at high confluency on 0.4 um PET transwell inserts (Corning 353090). Nine 6-well transwells were pooled for each replicate sample. Monolayers were differentiated for 7 days and Halo fusion proteins were induced with 0.5 µg/mL doxycycline. Following PBS wash, cells were incubated with 1 µM DBF Halo ligand in HBSS (gift from Rob Spitale) at 37°C for 15 minutes. Negative control samples lacking DBF were incubated in HBSS only.

Cells were washed with full growth media for 10 minutes at 37°C twice. Immediately following this, cells were incubated in 1mM propargylamine (Sigma-Aldrich P50900) in HBSS for 5 minutes at 37°C. Cells were then moved to the green light chamber containing two 100 W LED flood lights (USTELLAR) in an enclosed dark space. Cells were irradiated with green light for 10 minutes in the room temperature chamber sandwiched between two flood lights. Cells were collected in Trizol (Ambion) and homogenized by passing lysates through a 20G needle 20 times. Total RNA was then isolated following Trizol’s manufacturer instructions.

RNA was DNase treated using DNase I (NEB M0303S) for 30 minutes at 37°C. RNA was then recovered using Quick RNA columns (Zymo Research R1055) following the sample clean up protocol and eluted in 100 µL water.

### In vitro biotinylation of RNA via “Click” chemistry

Approximately 200 µg of total RNA was used in the Click reaction. The Click reaction contained 10 mM Tris pH 7.5, 2 mM biotin picolyl azide (Sigma-Aldrich 900912), 10 mM sodium ascorbate made fresh (Sigma 11140), 2 mM THPTA (Click Chemistry Tools 1010-100), and 100 µM copper sulfate (Fisher Scientific AC197722500). Click reactions were split into 50 µL reaction volumes and then incubated for 30 minutes in the dark at 25°C. Click reactions were pooled and cleaned using a Quick RNA Mini kit (Zymo Research R1055) following the sample clean up protocol and eluted in 100 µL water. Negative control samples where cells were not treated with DBF were also biotinylated using this procedure.

### Streptavidin pull down

100 µg of biotinylated RNA at 1 µg/µL was used as the input for streptavidin pulldowns. Different volumes of streptavidin-coated magnetic beads (Pierce PI88816) were used depending on the amount of labeled RNA by each Halo fusion protein. Halo-P65 required 1 µl beads per 2 µg total RNA while apical-Halo and basal-Halo required 1 µl beads per 4 µg RNA. Prior to mixing with RNA, beads were washed 3 times in B&W buffer (5 mM Tris pH 7.5, 0.5 mM EDTA, 1M NaCl, 0.1 Tween 20), 2 times in solution A (0.1 M NaOH, 50 mM NaCl), and 1 time in solution B (100 mM NaCl). The beads were resuspended with 100 µg RNA and 50 mM NaCl to a total volume of 100 µL. The pulldown reaction was rotated for 2 hrs at 4°C. The beads were then washed 3 times for 5 minutes each in B&W buffer with rotation at room temperature.

RNA was then eluted from the beads by resuspending in 50 µL PBS and 150 µL Trizol and incubated for 10 minutes at 37°C. The eluted RNA was recovered from this mixture using a DirectZol micro RNA columns (Zymo Research R2062) following the manufacturer’s instructions and eluted in 10 µL of water. The eluted RNA concentration was measured by Qubit high sensitivity RNA kit (Invitrogen Q32855) and +DBF and -DBF samples were compared.

### RNA dot blots

The efficiency of the biotinylation reaction and the streptavidin pulldown were assayed by RNA dot blots. Hybond-N+ membrane (GE) was wet with 2X SSC then allowed to dry for 15 minutes. 5 µg of biotinylated RNA samples or 1% of streptavidin eluted RNA were spotted on the membrane and allowed to dry for another 30 minutes. To stain for total RNA, the blot was incubated in 1% methylene blue (VWR 470301-814) for 2 minutes and destained using deionized water. Next, the membrane was blocked using 5% BSA for 30 minutes and washed 3 times in PBST (PBS + 0.01% Tween). Biotinylated RNA was then detected using streptavidin-HRP (Abcam ab7403) at a dilution of 1 to 20,000 in 3% BSA by addition to the membrane with rocking overnight at 4°C. Membranes were washed 3 times for 10 min each in PBST at room temperature. Streptavidin-HRP was detected using standard HRP chemiluminescent reagents (Advansta) and visualized using chemiluminescent imaging on a Sapphire molecular imager (Azure Biosystems).

### Imaging of alkynylated molecules in cells

Halo-transgene expressing C2bbe1 cells were plated at high confluency on poly-D-lysine-coated coverslips (neuVitro). Monolayers were differentiated for 7 days with 0.5 µg/mL Doxycycline to induce Halo fusion protein expression. Media was changed every 3-4 days. Cells were treated with DBF, propargylamine, and irradiated with green light exactly as described for Halo-seq. Instead of lysing cells with Trizol to recover RNA, cells were washed with PBS 3 times and fixed and permeabilized with 3.7% formaldehyde and 0.3% Triton-X (VWR 80503-490) in PBS for 30 minutes at room temperature. Following PBS wash, *in situ* click reactions were performed by incubating cells with 100 µL Click buffer (100 µM copper sulfate, 2 mM THPTA, 10 mM fresh sodium ascorbate, 10 µM Cy3 picolyl azide (Click Chemistry Tools 1178-1)) for 1 hour at 37°C in the dark. Two negative controls were included: samples with DBF omitted from the in-cell labeling reaction and cells with Cy3 picolyl azide omitted from the *in situ* Click reaction. Following the Click reaction, the coverslips were washed three times for 5 minutes each in wash buffer (0.1% Triton, 1 mg/mL BSA in PBS). Next, coverslips were incubated in wash buffer supplemented with 100 ng/mL DAPI for 30 minutes at 37°C and then washed twice with wash buffer for 5 minutes each. The coverslips were mounted with fluoromount G and imaged using a Deltavision Elite widefield fluorescence microscope (GE).

### Library preparation and sequencing

rRNA-depleted RNAseq libraries were prepared using an RNA HyperPrep Kit (KAPA / Roche). 100 ng of RNA was input into the procedure and fragmented for 3.5 minutes at 94°C. Libraries were amplified using 14 PCR cycles.

Libraries were sequenced using paired end sequencing (2 × 150 bp) on a NovaSeq high-throughput sequencer (Illumina) at the University of Colorado Genomics Core Resource. Between 20 and 40 million read pairs were sequenced for each sample.

### Computational strategy for HaloSeq

Library adapters were removed using cutadapt 2.1 (Martin, 2011). Transcript abundances in RNAseq data were quantified using Salmon 0.11.1 (Patro et al., 2017) and a human genome annotation retrieved from GENCODE (www.gencodegenes.org, GENCODE 28). Gene abundances were then calculated from these transcript abundances using tximport (Soneson et al., 2015), and genes whose abundance in input and streptavidin-pulldown samples were identified using DESeq2 (Love et al., 2014).

### Analysis of pre-mRNA transcripts in RNAseq data

To quantify the relative abundances of pre-mRNA (including introns) and mature mRNA (all introns spliced), a custom fasta file was supplied to Salmon which contained two versions of every transcript, one with all introns remaining and one with all introns removed. The custom fasta file was generated using this script: https://github.com/rnabioco/rnaroids/blob/master/src/add_primary_transcripts.py. Salmon then assigned reads competitively to these transcripts.

### Immunofluorescence staining in monolayers

C2bbe1 cells were seeded on PDL coated glass coverslips (neuVitro) within 12-well plates at high confluency. Monolayers were allowed to differentiate for 7 days, and media was changed every 3-4 days. Following PBS wash, cells were fixed with 3.7% formaldehyde at room temperature for 15 min. After another PBS wash, cells were permeabilized for 30 minutes at room temperature with 0.3% Triton-X (VWR 80503-490) in PBS. Primary antibodies with various dilutions (Ezrin, Cell Signaling Technology 3145S, 1:250; NaK ATPase, DSHB A5-s, 1:50; TOM20, ProteinTech 11802-1-AP, 1:250) were diluted in PBS containing 1% BSA and 0.3% Triton-X for 1 hr at 37°C. Cells were then rinsed three times with PBS before adding 100 ng/mL DAPI and fluorescent secondary antibody (Cell Signaling Technology 4413S) diluted 1:500 in PBS containing 1% BSA and 0.3% Triton-X for 1 hr at room temperature. When imaging for mitochondria, 100 ng/mL MitoView Green (Biotium 70054-T) is added at this point as well. Following PBS wash, cells were mounted onto slides with Fluoromount G (SouthernBiotech 0100-01) and sealed with nail polish. Slides were imaged on a widefield DeltaVision Microscope at 60X (GE) with consistent laser intensity and exposure times across samples.

### SmiFISH probe design and preparation

To assay RNA localization of endogenous genes, the R script Oligostan (Tsanov et al., 2016) was used to design primary probes for each RNA of interest. Probes were 26-32 nt in length with 40-60% GC content and repetitive sequences were avoided. Primary probes were designed with excess sequence for hybridization to a fluorescence FLAP sequence.

FLAP Y (TTACACTCGGACCTCGTCGACATGCATT) covalently attached to Cy3 was used for all smiFISH experiments targeting endogenous RNA. 1X pmol of Primary probes and 1.2X pmol FLAP sequences were hybridized in NEB buffer 3 in a thermocycler. The reaction was heated to 85°C for 3 minutes, cooled to 65°C for 3 minutes then held at 25°C yielding hybridized smiFISH probes specific to the endogenous RNA target.

### Stellaris probe design and preparation

Customized Stellaris smFISH probes were designed and ordered using the freely available probe designer software by Biosearch Technologies. Probes that were designed across the open reading frame of Firefly luciferase were used to assay localization of Firefly reporter constructs. Probes were resuspended in water to a final concentration of 12.5 µM.

### Cell preparation and hybridization for FISH experiments

Generally, FISH experiments were performed as previously described (Arora et al., 2022). CAD cells expressing reporter constructs were induced with 2 µg/mL doxycycline for 48 hours. Cells were then plated on PDL coated glass coverslips (neuVitro) within 12-well plates at approximately 2.5 × 104 cells per well in full growth media. Cells were allowed to attach for 2 hours before changing to serum depleted media supplemented with 2 µg/mL doxycycline for 48 hours.

C2bbe1 cells expressing reporter constructs were seeded on PDL coated glass coverslips at high density and allowed to differentiate into monolayers for 7 days in full growth media supplemented with 2 µg/mL doxycycline. Media was changed every 3-4 days. Cells were washed once with PBS before being fixed with 3.7% formaldehyde (Fisher Scientific) for 10 minutes at room temperature. Following 2 PBS washes, cells were permeabilized with 70% Ethanol (VWR) at 4°C for 6-8 hours or at room temperature for 2 hours. Cells were incubated in smFISH wash buffer (2x SSC with 10% formamide) at room temperature for 5 minutes. Per coverslip, 2 µL of Stellaris FISH Probes, or hybridized smiFISH probes were added to 200 µL of hybridization Buffer (10% dextran sulfate, 10% formamide in 2X SSC). Using a homemade hybridization chamber made from an empty tip box with wet paper towels and parafilm, coverslips were incubated cell side down in the hybridization buffer overnight at 37°C. Coverslips were washed with wash buffer in fresh 12-well plates cell side up for 30 minutes at 37°C in the dark. 100 ng/mL DAPI was diluted in wash buffer and added to the cells in the dark at 37°C for 30 minutes. Cells were then washed for 5 minutes at room temperature with wash buffer. Coverslips were then mounted onto slides with Fluoromount G and sealed with nail polish. Slides were imaged at 60X on a widefield DeltaVision Microscope with consistent laser intensity and exposure times across samples.

### FISH computational analysis

FISH-quant (Mueller et al., 2013) was used to identify smFISH puncta within CAD neurites and somas through use of its built-in Gaussian filtering and spot detection. All spot thresholding parameters were set to exclude gross outliers. The detection settings were the same for every soma and neurite imaged within an experiment. Approximately ten cells were analyzed for each sample. The mean number of Firefly luciferase transcripts in neurites compared to soma was calculated by FISH-quant mature mRNA summary files.

Images of epithelial monolayers were analyzed similarly with cell outlines approximated from GFP expression. Instead of using summary files, all detected spot files were used to ensure thresholding included spots from positive controls while excluding spots detected in negative control samples in which the RNA species (e.g. Firefly luciferase) being probed was not expressed.

Firefly luciferase reporter transcript puncta were thresholded until an expected expression of 100-1000 spots per cell was achieved. smiFISH puncta counts of different transcripts were expected to approximately reflect relative expression values calculated from RNA sequencing.

### Image analysis

Images were collected using a widefield Deltavision Elite microscope at 60X magnification. All images were deconvoluted by Deltavision’s default software, SoftWorx. Images of monolayers were collected as large stacks with many slices 0.2 µm apart from each other. FIJI(Schindelin et al., 2012) was used to filter all images in a stack using the default “rolling ball” algorithm before being average or max projected through the XY, XZ and YZ planes by the XYZ projection plugin (Omer et al., 2018). Average projections were used for all protein signal while max projections were used for smFISH signals. Projections were brightness and contrast matched for all relevant samples. RGB profile lines were drawn and analyzed with RGB profiler (Christophe Laummonerie, 2004).

CAD neurites were also imaged as stacks. Images were filtered using the default “rolling ball” algorithm and max projected through the XY.

### Neuronal fractionation sequencing computational strategy

Transcript abundances were calculated by salmon v0.11.1 (Patro et al., 2017) using Gencode genome annotations. Transcript abundances were then collapsed to gene abundances using txImport (Soneson et al., 2015). Localization ratios were calculated for each gene as the log2 of the ratio of neurite/soma normalized counts for a gene produced by DESeq2 using the salmon/tximport calculated counts (Love et al., 2014). Genes were required to have a minimum of 10 counts in any sample for analysis.

### siRNA depletion of Larp proteins

Mouse specific siRNAs were ordered from IDT for each Larp target. siRNAs were transfected into cells with Lipofectamine RNAiMAX (Thermo Fisher, 13778100). Briefly, 0.5 µL of 10 µM siRNA was combined with 50 µL Optimem media and 1.5 µL RNAiMAX reagent for roughly 1.0 × 105 cells in one well of a 24-well plate. CAD cells were transfected in the presence of doxycycline to induce expression of reporter constructs. Media was replaced 24 hours after transfection with full growth media supplemented with 2 µg/mL doxycycline. 24 hours following media change, cells were plated on PDL-coated coverslips (if imaging) and after attaching, media was changed again for serum free media with 2 µg/mL doxycycline to induce neurite outgrowth for 48 hours. Negative control siRNAs from IDT were used to control for effects due to transfection. For all proteins tested, knockdown of expression was confirmed by qPCR normalized to HPRT. For Larp1, immunoblots confirming protein knockdown were performed with mouse monoclonal Larp1 antibody (Santa Cruz sc-515873).

### qPCR of endogenous transcripts

SYBR green (Biorad) mastermix was used to analyze RNA knockdown of Larp siRNAs and LARP1 rescue in C2bbe1 cells. RNA was collected from siRNA transfected CAD cells as detailed above. Total RNA was collected from C2bbe1 monolayers plated at high confluency in 12 well plates. For C2bbe1 cells, media containing 2 µg/mL doxycycline was changed every 3-4 days for 7 days of differentiation. Following differentiation of both cell types, media was removed from cells and replaced with RNA Lysis Buffer (Zymo Research, R1060-1). Cells were lysed at room temperature for 15 minutes on a rocker. RNA was collected using Quick RNA micro columns (Zymo Research, R1051). The provided on-column DNAse treatment was performed for 15 minutes at room temperature. 500 ng of purified RNA was used to synthesize cDNA using iScript Reverse Transcriptase Supermix (BioRad, 1708841). cDNA was combined with gene specific primers and iTaq Universal SYBR green master mix (BioRad, 1725122). Gene expression of siRNA targeted Larps was normalized to HPRT before being compared to the expression in negative control siRNA targeted samples.

### Immunoblotting

Expression of Larp1 siRNA treated and CRISPR/Cas9 KO cell lines were determined by immunoblot analysis. Lysates were collected by scraping cells into 100 µL RIPA buffer, incubated on ice for 15 minutes then sonicated for 30 seconds. Lysates were stored at -80°C. Lysates were denatured with sample buffer (Invitrogen NP0008) and boiling at 100°C for 5-10 minutes. Denatured lysates were separated by PAGE on 4%–12% Bis-Tris gradient gels along with Flash Protein Ladder (Gel Company FPL-008) using MOPS SDS NuPAGE Running Buffer (Thermo Fisher, NP0001) at 200 V for 1 hour. Gels were transferred to PVDF membranes using an iBlot2 dry transfer device (Thermo Fisher, IB24002). Total protein was assayed by incubation with Ponceau staining (Sigma-Aldrich P7170), which was later destained and imaged on a Sapphire biomolecular imager (Azure Biosystems). Blots were then blocked with agitation in 5% milk powder (Sigma) in PBST (PBS + 0.01% Tween) for 1 hr at room temperature. Blots were washed 3 times, 5 minutes each in PBST at room temperature with agitation before being incubated in 1:100 LARP1 mouse monoclonal (Santa Cruz) primary antibody in 2% BSA (Research Products International A30075) in PBST overnight at 4°C with agitation or with 1:5000 beta actin mouse monoclonal antibody (Sigma A5441) in 2% milk powder in PBST for 2 hours at room temperature with agitation. Blots were washed 3 times, 5 minutes each at room temperature with agitation and then incubated with Anti-mouse IgG HRP-conjugated secondary antibody (Cell signaling 7076S) in 2% milk powder in PBST and incubated at room temperature for 2 hours. Blots were washed again in PBST and visualized using WesternBright HRP Substrate Kit (Advansta, K-12043-D10) and Sapphire biomolecular imager (Azure Biosystems) set to collect chemiluminescent signal.

### Morphology analysis of CAD cells

CAD cells were siRNA treated with various targeting or negative control siRNAs as detailed above. Differentiation occurred in a single well of a 6-well plate. An EVOS imager (Thermo Fisher) at 20X was utilized to take tiled brightfield images across the entire well. Images were analyzed in FIJI (Schindelin et al., 2012) using high contrast settings and NeuronJ (Meijering, 2004) was used to measure neurite lengths. The same images were used to count cells with and without neurites for qualitative morphological analyses.

## PUBLISHED DATA SETS USED

Moor 2017, Mouse LCM RNAseq (GSE95416: GSE95276; PRJNA377081)

Dermit 2020, projection localization RNA seq (E-MTAB-8470, E-MTAB-9520, and E-MTAB-9636) Cassella and Ephrussi 2021, Drosophila Follicular Cell LCM RNAseq (E-MTAB-9127)

## DATA AVAILABILITY

All high-throughput RNA sequencing data as well as transcript quantifications have been deposited at the Gene Expression Omnibus under accession number GSE200004.

## SUPPLEMENTARY TABLES

**Table S1**. Localization values in C2bbe1 monolayers. DESeq2 output of Cytoplasmic Bias (CB) and calculated FDR values (CB_FDR). Cytoplasmic bias is the log2 of a gene’s cytoplasmic enriched abundance divided by the abundance in total RNA. Apical Bias (AB) and AB_FDR values are also included for all observed genes where Apical bias is the log2 of a gene’s apical enriched abundance divided by its basal enriched abundance.

## SUPPLEMENTARY FIGURES

**Figure S1.**
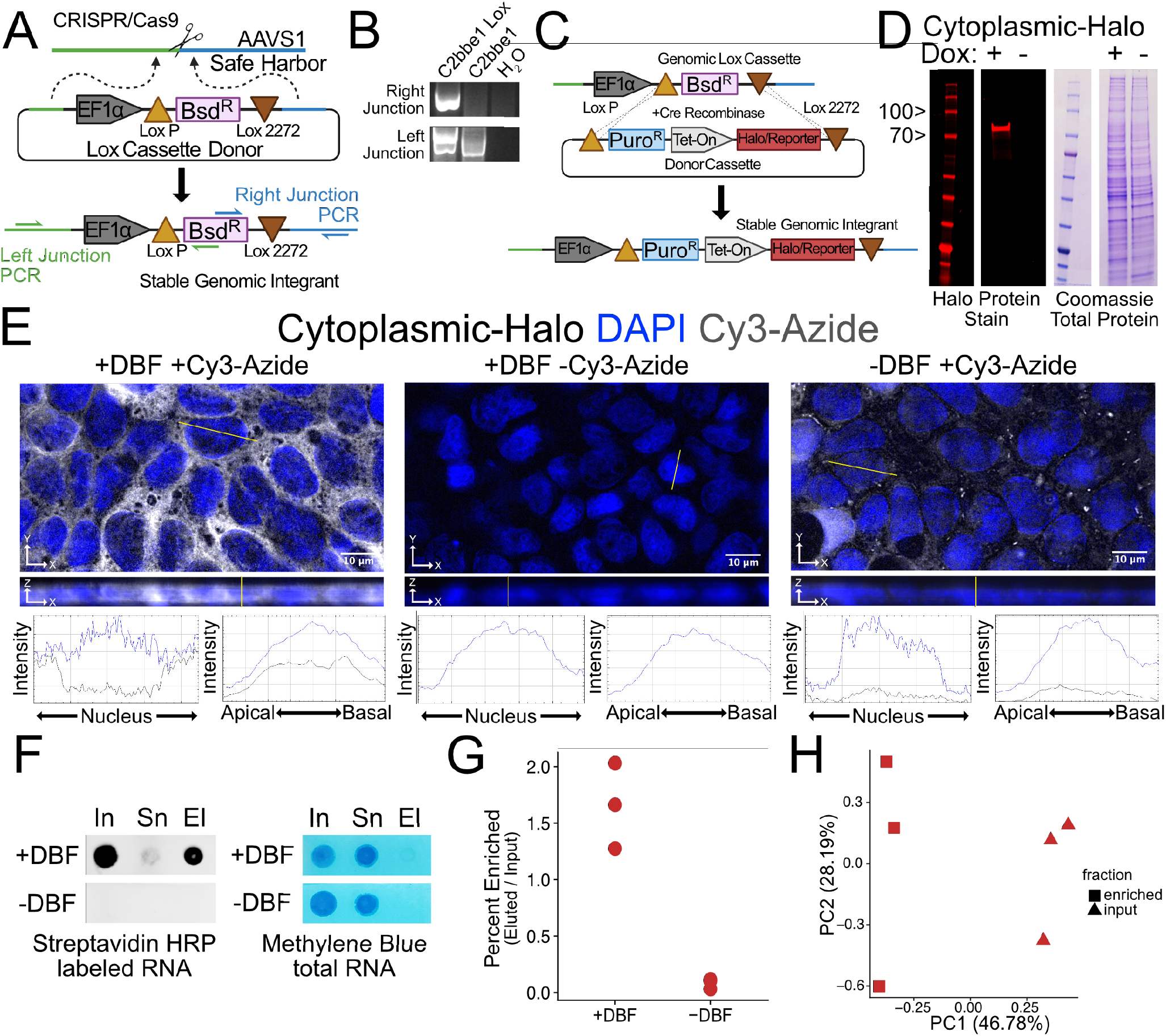
Halo-seq requires transgene expression to enrich cytoplasmic RNAs via biotinylation. (A) Schematic of Lox cassette knock-in to the AAVS1 safe harbor locus. A loxP-flanked blasticidin resistance gene driven off a constitutively active EF1α promoter is inserted into the AAVS1 safe harbor. Primer positions to PCR amplify the junctions of this insertion are notated. (B) PCR amplification of knock-in junctions. A cell line without the knocked in cassette and water included as template controls. (C) Schematic of cre-mediated cassette switching. Through cre recombinase activity, the floxed blasticidin gene is replaced with a puromycin resistance gene as well as any additional reporter or HaloTag fusion proteins. Stable cassette switching events are then selected with puromycin to produce stable cell lines. (D) SDS-PAGE gel of doxycycline-induced cytoplasmic-Halo construct (95 kDa) visualized with Halo ligand fluorophore in red. Total protein is visualized with a Coomassie stain. (E) Fusing Cy3-Azide fluorophores to alkynylated biomolecules using Click chemistry allows for visualization of labeled molecules *in situ*. Alkynylated molecules are restricted to the cytoplasm in cells expressing cytoplasmic-Halo. This localization is dependent on both addition of DBF and Cy3-azide, demonstrating the ability of HaloTag-restricted DBF to induce alkynylation of biomolecules specifically in the cytoplasm. (F) *in vitro* biotinylation of alkynyklated RNA labeled from cytoplasmic-Halo expressing cells is dependent on DBF addition as visualized by streptavidin-HRP on an RNA dot blot. Biotinylated input (In) RNA are cleared from the supernatant (Sn) by streptavidin pull-downs but efficiently eluted (El) off the beads. Methylene Blue stains total RNA. (G) Percent enriched RNA after streptavidin pull-down of cytoplasmic-Halo labeled RNAs. (H) Principal component analysis of gene expression values from Halo-seq in cytoplasmic-Halo cells.

**Figure S2.**
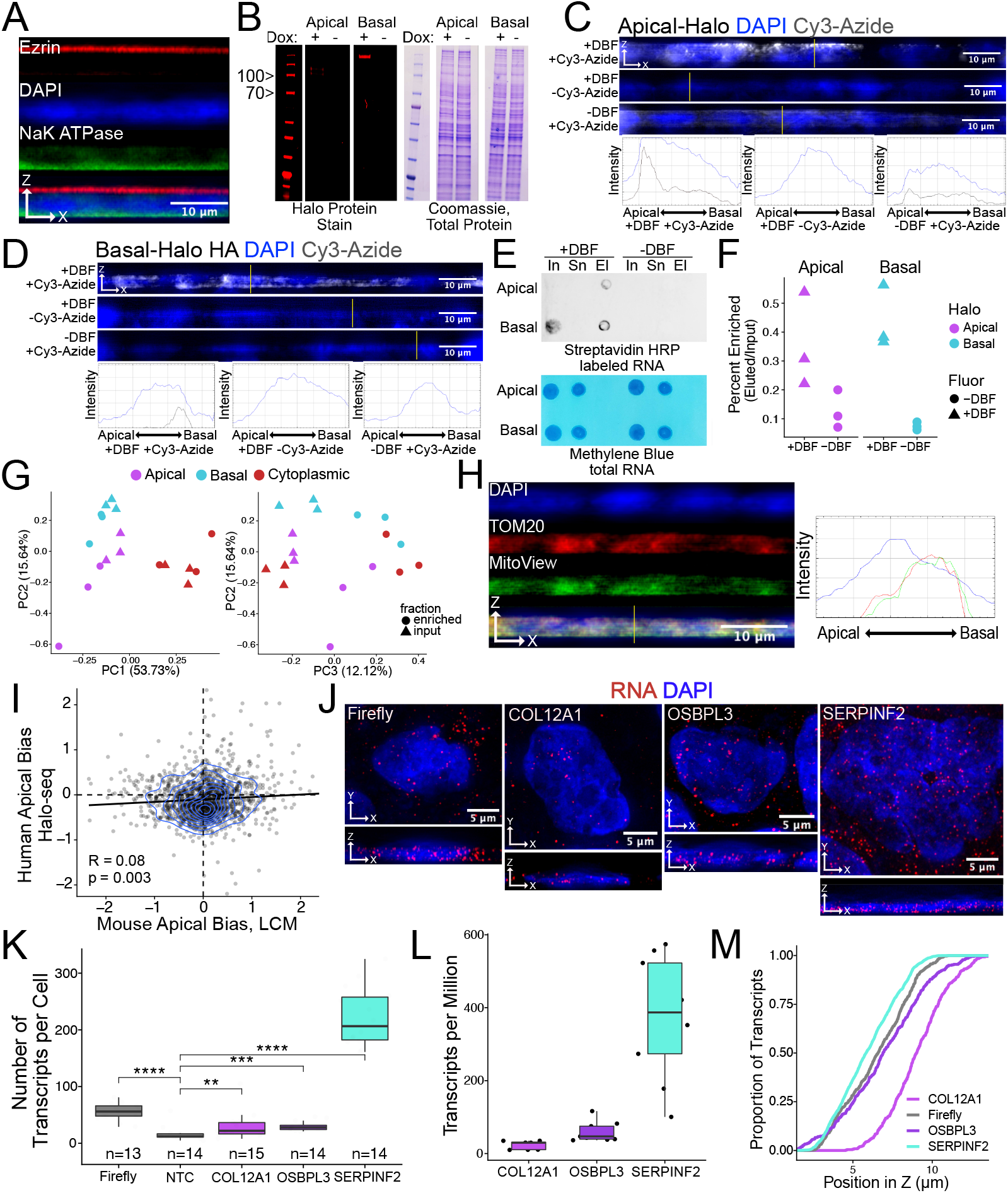
Halo-seq enriches RNA localized across the apicobasal axis via biotinylation. (A) C2bbe1 monolayers differentiated for 7 days on transwell inserts are highly polarized as demonstrated by immunofluorescence for Ezrin (apical) and NaK-ATPase (Basal) endogenous polarity protein markers. Images are an average projection through the XZ axis of many cells. (B) SDS-PAGE gel of doxycycline-induced apical-Halo and basal-Halo constructs (95 and 150 kDa respectively) visualized with Halo ligand fluorophore in red. Total protein is visualized with a Coomassie stain. (C) Cy3-Azide fluorophores attached to alkynylated biomolecules by Click chemistry visualize labeled molecules *in situ*. Alkynylated molecules are restricted to the apical or (D) basal pole in cells containing apical-Halo and basal-Halo respectively. This localization is dependent on both addition of DBF and Cy3-azide. Images are an average projection through the XZ axis of many cells. (E) *in vitro* biotinylation of RNA labeled in apical-Halo and basal-Halo expressing cells is dependent on DBF addition as visualized by streptavidin-HRP on an RNA dot blot. Biotinylated input (In) RNA are cleared from the supernatant (Sn) by streptavidin pull-downs but efficiently eluted (El) off the beads. Methylene Blue stains total RNA. (F) Percent enriched RNA after pull-down of apical-Halo and basal-Halo labeled RNAs. (G) Principal component analysis of gene expression values from the cytoplasmic-Halo, apical-Halo and basal-Halo experiments. PC1 and 2 separate Halo-construct lines while PC2 and 3 separate input and enriched fractions. (H) Mitochondria are visualized with both MitoView Green and immunofluorescence for TOM20 protein. Profile lines across the XZ axis show basal localization of mitochondria. Images are an average projection through the XZ axis of many cells. (I) Direct comparison of Halo-seq calculated Apical Bias in Human C2bbe1 monolayers and Laser Capture Microdissection (LCM) calculated Apical Bias in adult Mouse enterocytes (Moor et al., 2017). (J) Representative smiFISH images. Images are a max projection through the XZ axis of a single cell. (K) Number of puncta per cell. P-values were calculated using a Wilcoxon rank-sum test. No template control (Firefly luciferase smFISH probes in cells not expressing the Firefly luciferase transgene) included to show noise level in smiFISH. (L) Transcripts per million for each candidate localized RNA in each sequenced input sample (n=9). (M) Cumulative distribution of smiFISH spots across their position in Z as calculated by FISH-quant. P-values were all significant (< 0.05) compared to Firefly luciferase as calculated by Wilcoxon rank-sum test and printed on Figure 2G. ns (not significant) represents p > 0.05, * p < 0.05, ** p < 0.01, *** p < 0.001 and **** represents p < 0.0001.

**Figure S3.**
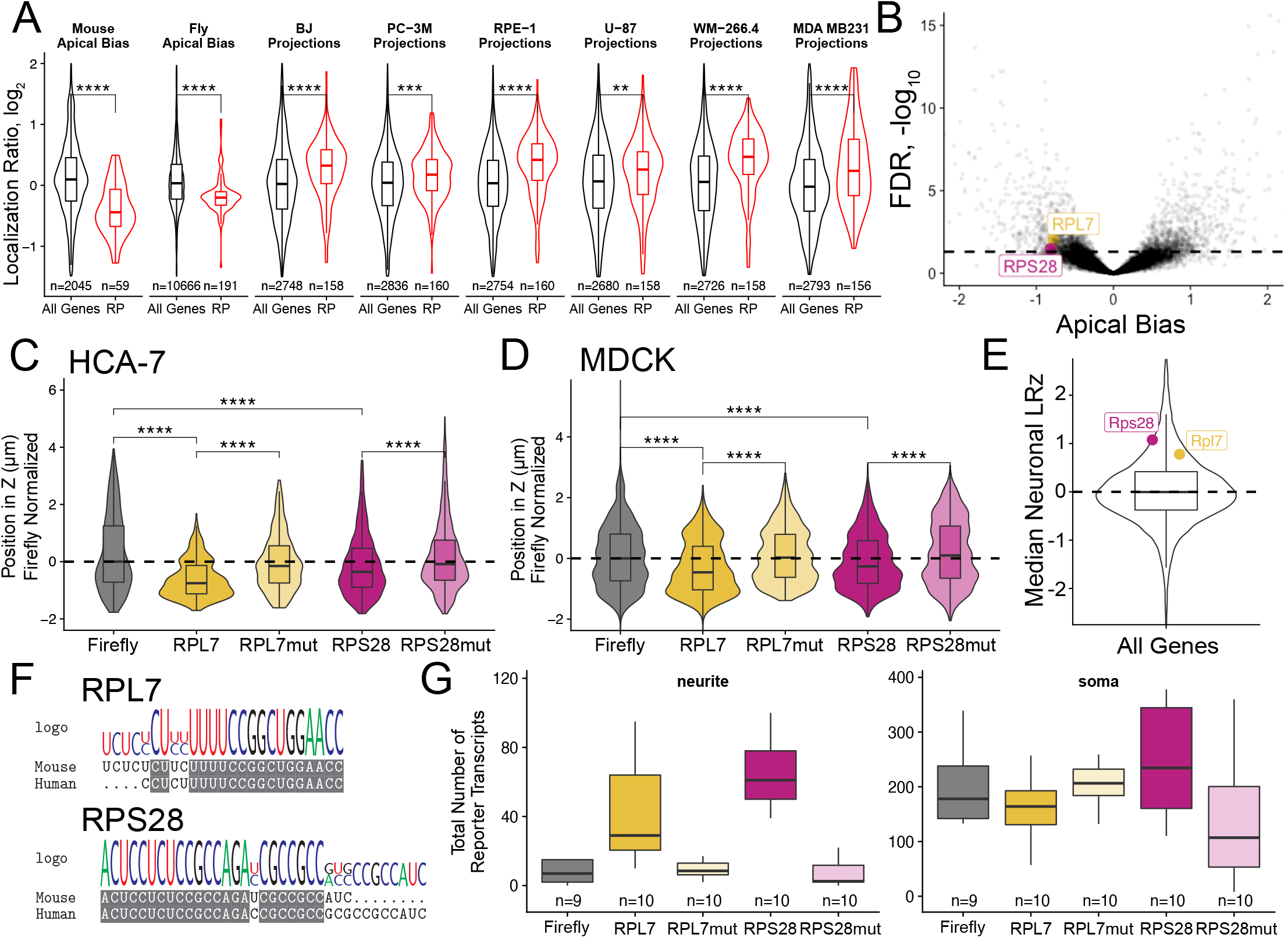
5′ TOP motifs are sufficient for RNA localization (A) Localization of ribosomal protein mRNAs across the apicobasal axis of adult mouse enterocytes (Moor et al., 2017), the apicobasal axis of drosophila follicular cells (Cassella and Ephrussi, 2021) and in migrating cell projections (Dermit et al., 2020) according to various sequencing experiments. P-values were calculated using a Wilcoxon rank-sum test. (B) Genes with differing Apical Bias with candidate RP mRNAs highlighted. P-values calculated by DESeq2 (C) 5′ TOP reporter transcript localization in HCA-7 monolayers. Reporter RNA smFISH spot position in Z normalized to median untagged Firefly luciferase transcript position. P-values were calculated using a Wilcoxon rank-sum test. (D) 5′ TOP reporter construct localization in MDCK monolayers. Reporter RNA smFISH spot position in Z normalized to median untagged Firefly transcript position. P-values were calculated using a Wilcoxon rank-sum test. (E) Neuronal localization ratio of all genes with candidate RP mRNAs highlighted. (F) Human and mouse 5′ UTR alignments for RPL7 and RPS28. (G) Total smFISH reporter construct puncta detected in neurites and soma by FISH-quant. ns (not significant) represents p > 0.05, * p < 0.05, ** p < 0.01, *** p < 0.001 and **** represents p < 0.0001.

**Figure S4.**
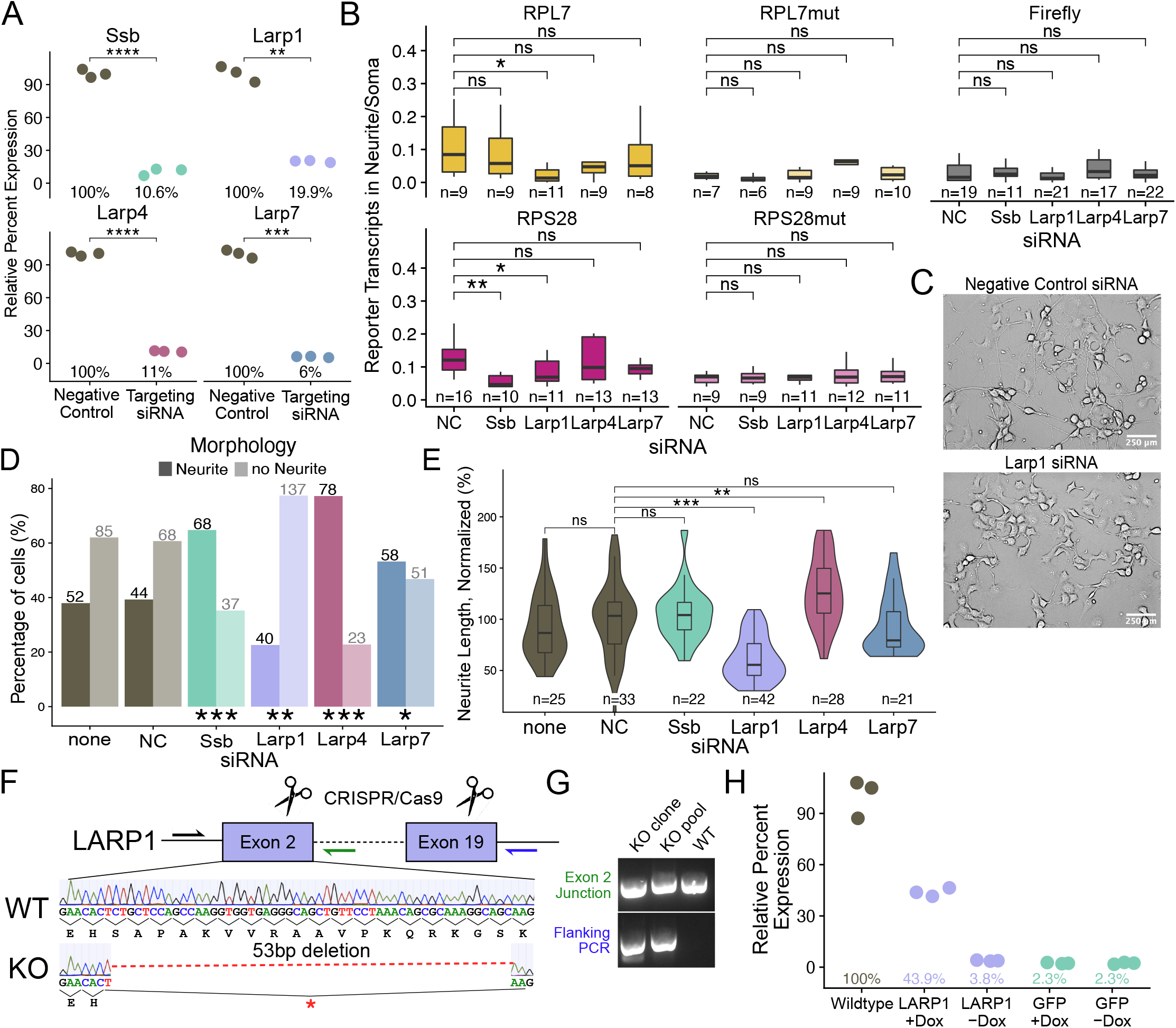
LARP1 mediates 5′ TOP localization to multiple cell morphologies. (A) Percent knockdown of Larp RNAs by targeting and negative control siRNAs as measured by qPCR normalized to HPRT. P-values were calculated using a student’s t test. (B) Reporter localization in neurons treated with different Larp siRNAs. Reporter RNA smFISH puncta in neurites normalized to soma. P-values were calculated using a Wilcoxon rank-sum test. (C) Representative brightfield images of Larp1 and negative control siRNA treated differentiated CAD cells (D) Proportion of cells with or without defined neurites when treated with different Larp siRNAs. Raw count data is printed. P-values were calculated using a Fisher’s exact test. (E) Neurite length of multiple Larp siRNA treated cells normalized to negative control (NC) siRNA treated cells. P-values were calculated using a student’s t test. (F) LARP1 CRISPR/Cas9 knockout Strategy. Guide RNAs targeting exon 2 and exon 19 facilitated loss of the intervening sequence and frameshifting deletions in Exon 2. Traces from sanger sequencing show a 53 base pair deletion in exon 2 that creates an early stop codon. (G) PCR amplification of the locus using primers that flank both gRNA cut sites shows loss of the sequence between exons 2 and 19. Exon 2 PCR shows decreased size due to deletions. (H) Relative expression of LARP1 rescue lines as normalized to HPRT then normalized to wildtype C2bbe1 LARP1 expression. ns (not significant) represents p > 0.05, * p < 0.05, ** p < 0.01, *** p < 0.001 and **** represents p <0.0001.

**Figure S5.**
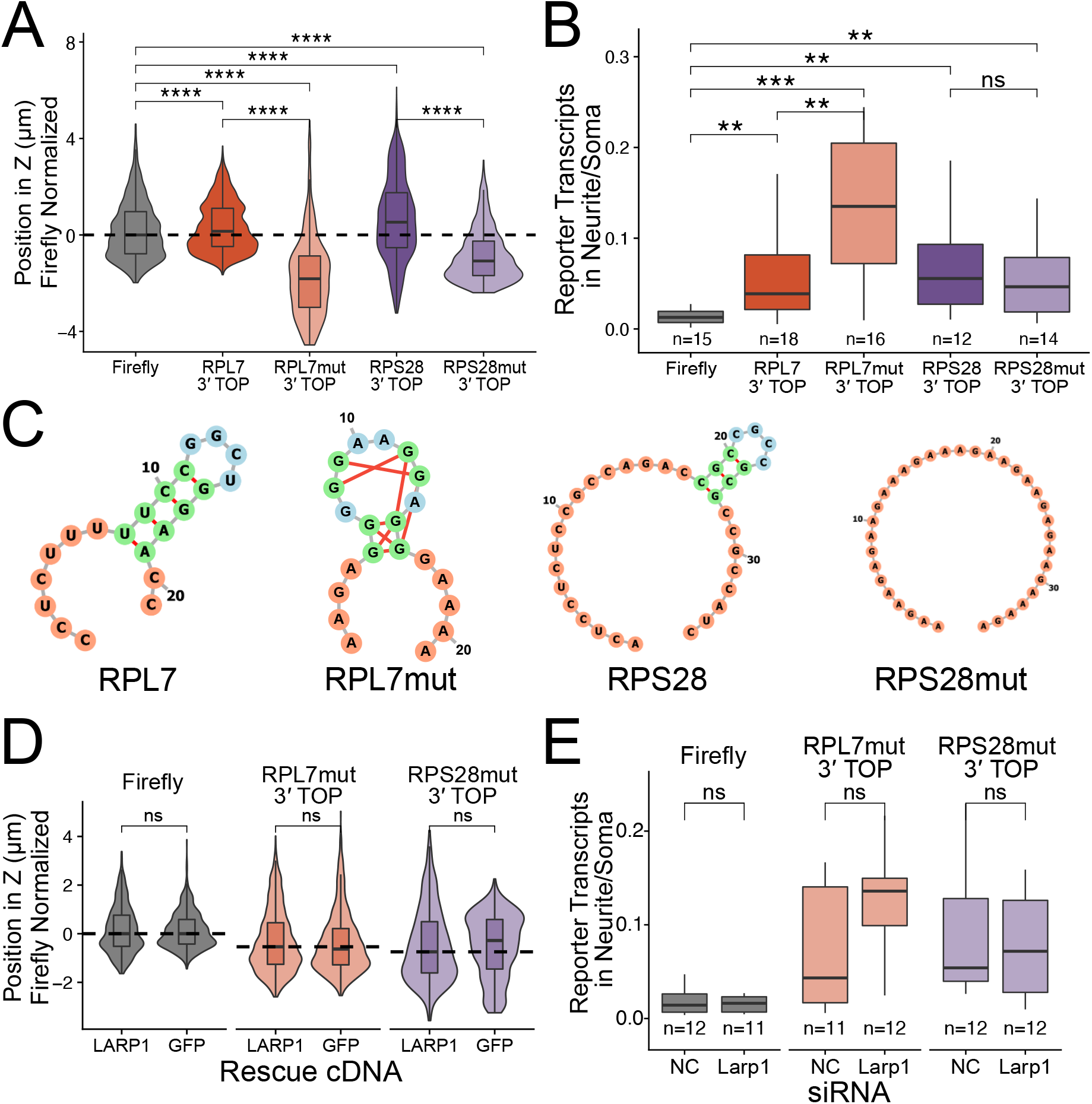
Moving the TOP motif away from the 5′ end of a transcript abolishes its RNA localization activity. (A) 3′ TOP reporter transcript localization in C2bbe1 monolayers. Reporter RNA smFISH spot position in Z normalized to median untagged Firefly luciferase transcript position. P-values were calculated by Wilcoxon rank-sum test. (B) Number of smFISH reporter transcript puncta detected in neurites normalized to soma. (C) TOP motif structural conformations as calculated by RNAfold (Lorenz et al., 2011). (D) 3′ TOP reporter localization in LARP1-depleted C2bbe1 monolayers. Reporter RNA smFISH puncta position in Z normalized to median untagged Firefly luciferase transcript position. P-values were calculated using a Wilcoxon rank-sum test. (E) Number of 3′ TOP reporter transcript puncta detected in neurites normalized to soma in Larp1 siRNA treated neurons. P-values were calculated using a Wilcoxon rank-sum test. ns (not significant) represents p > 0.05, * p < 0.05, ** p < 0.01, *** p < 0.001 and **** represents p < 0.0001.

**Figure S6.**
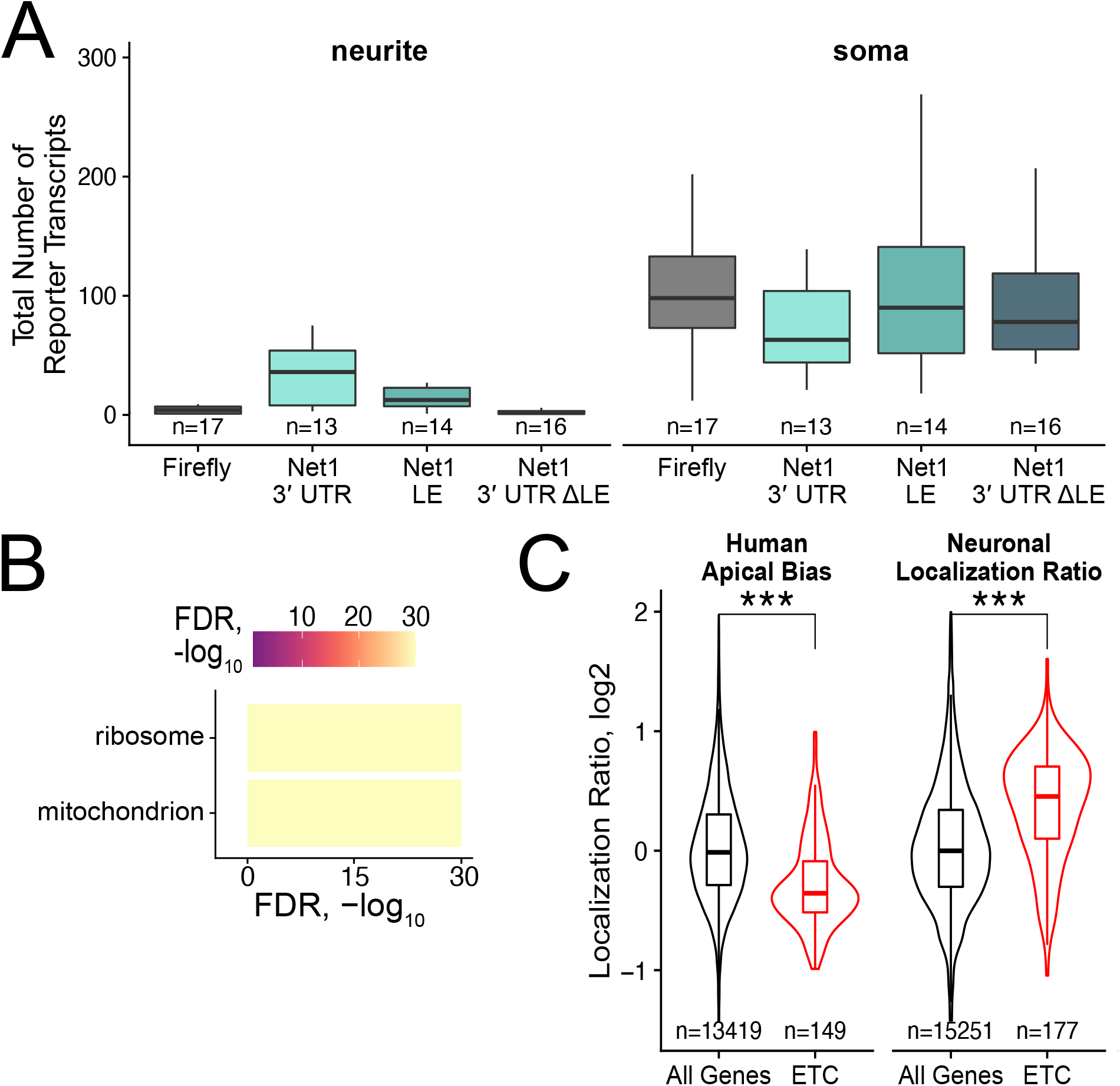
Organelle localization can explain RNA localization in different cell morphologies. (A) Total smFISH reporter transcript puncta detected in neurites and soma by FISH-quant. (B) Enriched gene ontology terms derived from RNAs identified as localized to the basal pole of enterocytes and to neurites of neurons. FDR calculated by topGO. (C) Localization of transcripts encoding electron transport chain (ETC) associated proteins in apicobasal and neuronal localization datasets. P-values were calculated using a Wilcoxon rank-sum test. ns (not significant) represents p > 0.05, * p < 0.05, ** p < 0.01, *** p < 0.001 and **** represents p < 0.0001.

**Figure S7.**
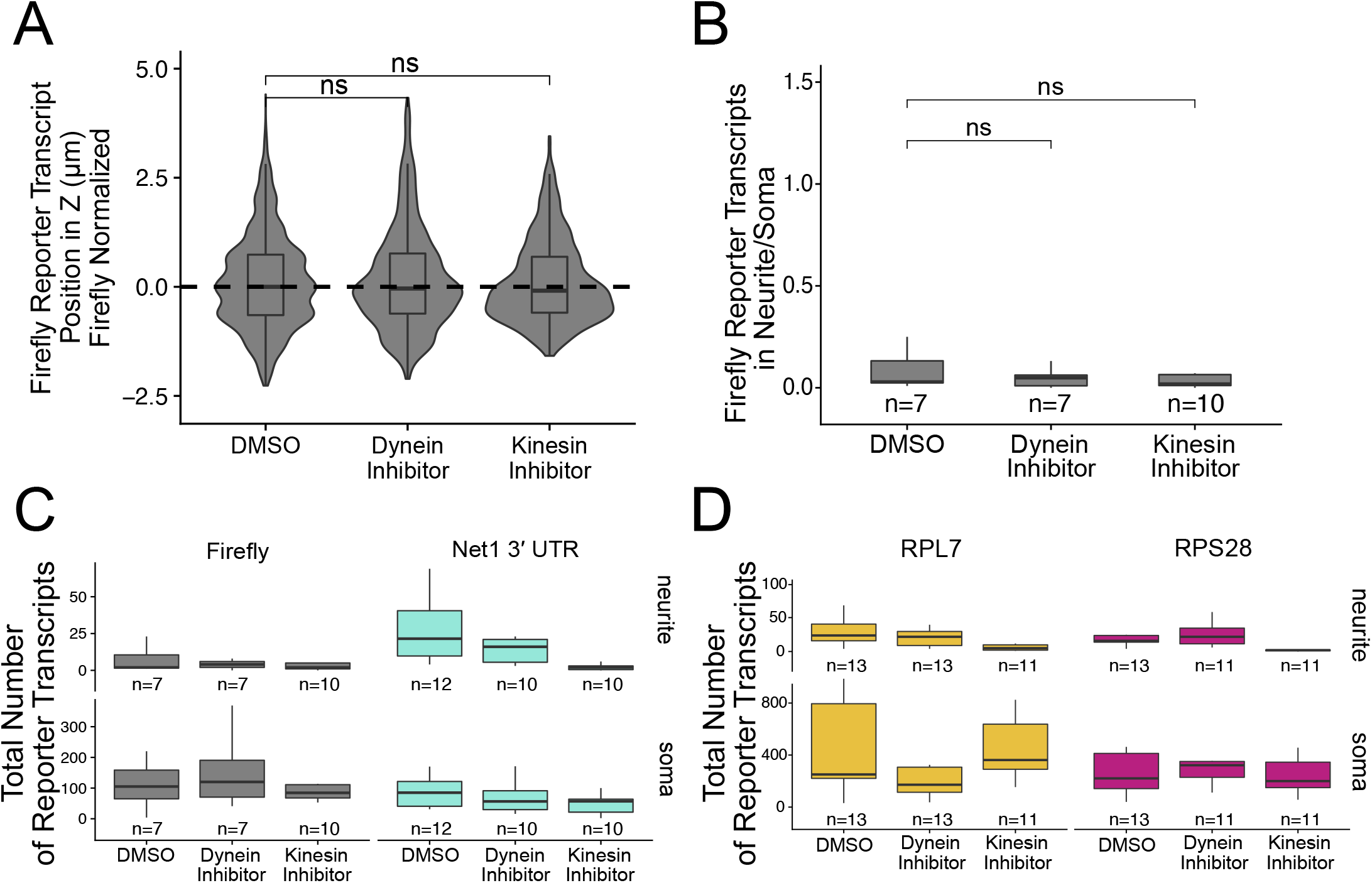
RNA localization requires kinesin in both neurites and enterocytes. (A) Firefly luciferase-Net1 3′ UTR reporter transcript localization with different drug treatments normalized to median untagged Firefly luciferase transcript position in Z. P-values calculated by Wilcoxon rank-sum test. (B) Number or Firefly luciferase transcripts detected in neurites normalized to soma with different drug treatments. P-values calculated by Wilcoxon rank-sum test. (C) Total number of Net1 and Firefly reporter transcript puncta detected in neurites and soma. (D) Total number of RPL7 and RPS28 5′ TOP reporter transcript puncta detected in neurites and soma. ns (not significant) represents p > 0.05, * p < 0.05, ** p < 0.01, *** p < 0.001 and **** represents p < 0.0001.

